# Domestication and deep lineage divergence define two discrete trajectories in *Kluyveromyces marxianus*

**DOI:** 10.64898/2026.04.28.721375

**Authors:** Jesús Martín Moreno-Hernández, Jolien Smets, Ana Lilia Torres-Machorro, Luis F. García-Ortega, J. Abraham Avelar-Rivas, Raúl A. Ortiz- Merino, Manuel R. Kirchmayr, Anna Muszewska, Lucia Morales, Kevin Verstrepen, Alexander DeLuna, Eugenio Mancera

## Abstract

Although genomics has greatly improved our ability to detect population structure and genetic divergence, it remains difficult to determine when diversity within a recognized species reflects continuous variation and when it marks the emergence of discrete evolutionary lineages. In yeasts, population-genomic studies often reveal overlapping signatures of environmental association, admixture, and domestication, making this distinction especially challenging. By analyzing 178 globally sampled genomes from *Kluyveromyces marxianus*, an emerging yeast of biotechnological interest, we show that diversity within this species resolves into two contrasting evolutionary trajectories that do not map onto broad geographic origin. Most industrial and clinical isolates belong to a lineage marked by gene loss, aneuploidy, and reduced fertility, consistent with genomic hallmarks of domestication. In contrast, isolates confined to traditional agave fermentations in Mexico and South Africa form a highly distinct lineage, separated from the other lineages by up to 4.8% genome-wide nucleotide divergence, widespread reciprocal monophyly, no detectable recent gene flow, and reduced mating compatibility despite co-occurrence with other lineages. Together, these results show that within a single yeast species, domestication-associated genome remodeling and deep lineage divergence can define distinct trajectories, with implications for strain selection in biotechnology.

## Introduction

Genomic data have transformed how biologists identify evolutionary lineages and delimit species, but they have also shown that genome-wide structure does not necessarily map onto reproductive isolation, ecological divergence, or formal taxonomy [1–5]. The problem is particularly evident within recognized species, where geographic structure, environmental specialization, admixture, and domestication can generate overlapping patterns of differentiation [6–9]. In such cases, a central question is whether diversity is best understood as a single structured continuum or as a set of more discrete evolutionary lineages. This distinction matters because it changes how discontinuities are interpreted, either as localized differentiation within an otherwise connected species or as evidence of more discrete lineage partitioning.

Yeasts offer a powerful system for investigating these dynamics. Their compact genomes, broad habitat ranges, recurrent association with human-managed environments, and expanding population-genomic resources have made them increasingly tractable models in ecology and evolution [10–13]. In yeasts, diversification can proceed through admixture, introgression, ploidy shifts, structural variation, and lineage splitting [12,14–16]. A well-studied example comes from the baker’s yeast *Saccharomyces cerevisiae*, where domesticated and wild populations differ in their dominant modes of genome evolution. Domesticated lineages are often enriched for ploidy variation, aneuploidy, genome-content change, and altered reproductive capacity, whereas wild populations are more strongly structured by single-nucleotide divergence and biogeography [12,14,15,17–19]. Even within bread-associated *S. cerevisiae*, industrial and artisanal strains appear to have followed distinct domestication trajectories [20]. Likewise, the dairy-associated yeast *Kluyveromyces lactis* includes a domesticated lineage with reduced diversity and introgressed lactose-utilization genes, suggesting that domestication-associated genomic divergence extends beyond *Saccharomyces* [21,22]. Yet in these systems, domestication-associated divergence remains embedded within a broader continuum of genomic variation. Whether this kind of genome remodeling can coexist with lineage divergence marked primarily by substantial nucleotide differentiation and limited gene flow remains less clear.

*Kluyveromyces marxianus* is an especially attractive system in this context. It is a fast-growing, thermotolerant non-conventional yeast with broad biotechnological relevance, including food fermentations, biomass conversion, and emerging biocatalytic applications [23–29]. At the same time, it is not restricted to industrial settings, but occurs across dairy, agave-associated, and other non-dairy environments [30–32]. This combination of applied relevance and ecological breadth makes *K. marxianus* a powerful system for examining the coexistence of linages with contrasting evolutionary trajectories. However, its population genomics remains far less resolved than that of *S. cerevisiae*. A first comparative survey based on a limited set of 14 available genomes identified three main phylogenetic groups associated broadly with dairy versus non-dairy origins, substantial variation in haplotype composition and ploidy, and a single highly divergent agave-associated isolate from South Africa [30]. A small number of additional genomes have since modestly expanded the known diversity of the species [31–34], but they have not yet clarified whether these discontinuities reflect sparse sampling along an otherwise continuous pattern of within-species variation or genuine partitioning into distinct evolutionary lineages.

Here, we analyzed 178 genomes of *K. marxianus* sampled globally from industrial, natural, clinical, and agave-associated sources to examine the observed discontinuities. We show that divergence in this species follows two contrasting genomic trajectories, with most industrial and clinical strains defining a domesticated lineage marked by gene loss, aneuploidy, and reduced fertility. In contrast, an agave-associated clade from Mexico and South Africa represents a case of deep lineage divergence, marked by pronounced nucleotide differentiation, limited gene flow, and reduced inter-lineage mating compatibility. Together, our results establish *K. marxianus* as a natural system for studying alternative routes of yeast diversification and provide a genomic framework for strain selection in biotechnology.

## Results

### *K. marxianus* is structured into three major lineages that do not map onto broad geography

Despite its recognized biotechnological relevance, the global population structure of *K. marxianus* remains poorly resolved. To establish a population-genomic framework for the species, we compiled and analyzed a dataset of 178 *K. marxianus* genomes, the vast majority newly sequenced for this study, from a broad range of natural and anthropogenic environments (**Figure 1**). This collection included dairy-associated industrial strains (25) and two from traditional beer production, 104 strains from agave-spirit production in traditional distilleries across Mexico [35], strains from natural sources, including soil, water, and overripe fruits (18), isolates from clinical cases involving livestock as well as human patients with mastitis and tuberculosis (8), and 21 strains from global culture collections with unknown isolation origin, including several patent-protected isolates. Although our collection was enriched for isolates from agave fermentations, the sampling was not narrowly local, as these fermentations are distributed across a broad region of Mexico, are also represented in South Africa, and have been shown to capture substantial genetic diversity for other yeast species [36,37]. Overall, the dataset spans multiple regions, with most isolates from North America and Europe and additional strains from South America, Africa, and Asia (**Figure 1a**). Full strain metadata and genomic features are provided in **Table S1**.

**Figure 1.**
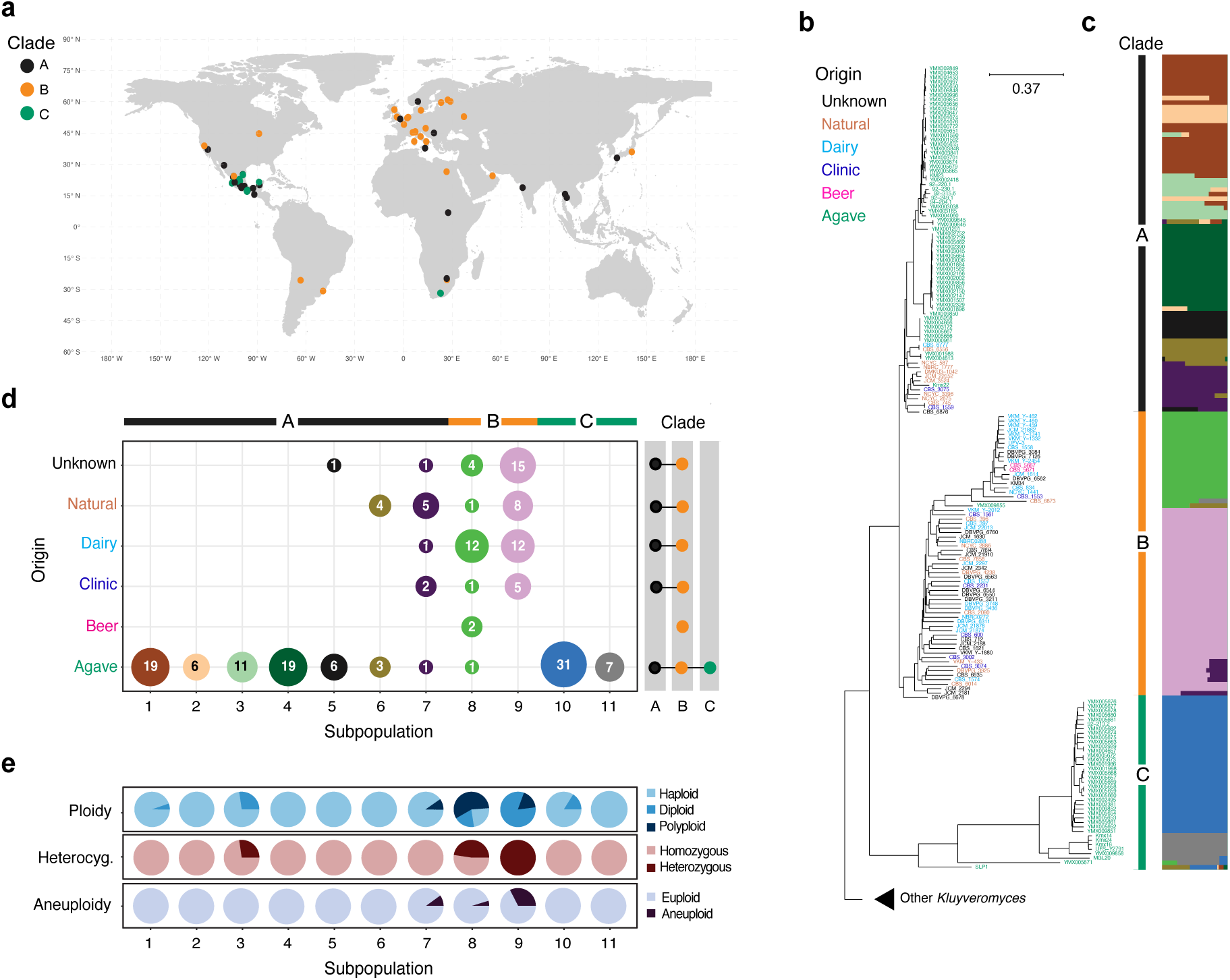
Global population structure of *K. marxianus* reveals three major lineages and eleven subpopulations. **a)** Global geographic distribution of the 178 strains. The color of the dots represents the phylogenetic clade to which each strain belongs, as in panel b. **b)** Neighbor-joining tree inferred from 953,880 biallelic informative markers. As a collapsed outgroup, we included genomic sequences of other *Kluyveromyces* species representing terrestrial (*K. lactis*, *K. dobzhanskii*, *K. wickerhamii*, *K. starmeri*) and aquatic (*K. aestuarii*, *K. siamensis*, *K. nonfermentans*) lineages. The color of the strain names at the tips represents the source of isolation as specified in the legend. **c)** Admixture analysis resolves the genetic diversity of *K. marxianus* (*n*=178) into eleven subpopulations (horizontal colored bars). K=11 was the value with the lowest cross-validation error (Figure S1); strains are sorted according to their phylogenetic position in panel b, and the vertical bar at the left in gray, orange, and green indicates the three major phylogenetic clades, A, B, and C, respectively. **d)** Co-occurrence of subpopulations (left panel) and clades (right panel) in each habitat of isolation. Numbers in the circles of the left panel represent the number of strains in each subpopulation and habitat. The colors of the circles denote the subpopulation as defined in panel c. **e)** Ploidy, heterozygosity, and aneuploidy across the eleven subpopulations of *K. marxianus*. Pie charts display the proportion of strains within each subpopulation exhibiting variation as indicated in the legend.

Whole-genome sequencing and analysis identified a total of 1,345,626 raw variants, most of them single-nucleotide polymorphisms (SNPs, 79.4%). The remaining variants were mostly short insertions and deletions (INDELs). The overall nucleotide diversity of the species was consistent with previous estimates (π = 1.5×10^-2^; [30]). This high average was largely driven by the diversity of strains from agave fermentations, which showed nearly the same value as the whole species. In contrast, dairy-industrial strains showed a twofold reduction (π = 7.0×10^-3^), and beer strains displayed the lowest diversity (π = 1.0×10^-3^), nearly an order of magnitude lower than agave-associated strains (**Figure S1**). Compared to other yeasts, *K. marxianus* is more diverse than *S. cerevisiae* (π = 3×10^-3^;[12]), but less diverse than its sister species *K. lactis* (π = 2.8×10^-2^; [22].

Using 953,880 biallelic informative sites, we inferred phylogenetic relationships among strains with a neighbor-joining tree [38], recovering three clearly separated lineages (**Figure 1b**). The first lineage, here termed clade A, comprised strains isolated from natural environments and agave fermentations. The second lineage, clade B, grouped most strains used in dairy and beer fermentations, together with a small number of clinical isolates. The third lineage, clade C, consisted exclusively of strains from agave fermentations. This last group included isolates recovered not only from fermentation tanks, but also from other substrates within the distilleries. Clades A and B were relatively close in the tree, whereas clade C was clearly more divergent from both. SNP density within clades followed the same pattern, with mean SNP densities of 8, 22, and 42 SNPs per kilobase in clades A, B, and C, respectively. Importantly, strains from clades A and B were broadly distributed across all four continents represented in our collection, whereas clade C strains were recovered exclusively from agave-based distilleries in Mexico and South Africa.

To further resolve population structure within these three lineages, we performed Admixture analysis. Cross-validation identified eleven subpopulations as the best-fit model (**Figure S2**). Clade A encompassed seven genetic clusters, while clades B and C were each split into two subpopulations (**Figure 1c**). The analysis also revealed ancestry sharing, with around 13% of strains showing mosaic patterns, particularly between subpopulations of clades A and B. This is consistent with previous work suggesting that diploid dairy strains from clade B may have originated from inter-lineage crosses with isolates from clade A [30]. Despite the large number of strains sequenced from clade C, evidence of admixture with clades A and B was detected in only two strains (YMX005671 and SLP1). Pairwise *F_ST_* values and principal component analysis (PCA) supported this pattern. Three subpopulations from clade A (1, 2, and 3) were among the least differentiated, with *F_ST_* values below 0.3, consistent with either more recent common ancestry or ongoing gene flow. In contrast, populations 10 and 11 within clade C displayed the highest *F_ST_* scores (∼0.9), indicative of strong differentiation and limited gene flow. In PCA, PC1 separated the compact cluster of clade A subpopulations from strains associated with anthropogenic niches, whereas PC2 distinguished the highly divergent clade C from all other genetic clusters (**Figure S2**).

To evaluate whether this lineage structure reflects broad geographic origin, we compared genetic clustering with the distribution of strains across habitats and regions. Most of the co-occurring subpopulation diversity was concentrated within the agave fermentation environment, where 90% (10/11) of the species subpopulations coexisted, despite originating from geographically distant locations. Unlike other related ascomycetous yeasts such as S*. cerevisiae* [12] and *K. lactis* [22], the population structure of *K. marxianus* does not appear to be organized primarily by broad geographic origin (**Figure 1d**). For example, individuals from lineage B were distributed across diverse locations, yet their genetic diversity was captured by only two subpopulations. In contrast, agave fermentations in Mexico harbored strains from ten of the eleven total subpopulations, including members of both clade A and clade C coexisting within the same fermentations despite strong genetic differentiation. A Mantel test using Mexican strains, the only subset with precise geographic origin for many isolates, detected a statistically significant correlation only within clade C, and even there the effect size was modest (*p* = 1×10^-5^, *r* = 0.56; **Figure S3**). Together, these results indicate that the major lineage structure of *K. marxianus* is only weakly associated with broad geographic origin.

The weak association between lineage structure and geography also provides context for the inferred phylogenetic placement of clade C. In rooted trees including other *Kluyveromyces* species as outgroups, clade C occupied the earliest-diverging position within *K. marxianus* (**Figure 1b**). Together with the limited admixture detected between clade C and the other major lineages (**Figure 1c**), this pattern suggests that clade C is unlikely to represent a recently derived subpopulation of the species. Instead, it supports clade C as an early-branching lineage within *K. marxianus*.

Beyond nucleotide divergence and ancestry patterns, the three major lineages also differed in ploidy and heterozygosity. Combining flow cytometry with *in silico* ploidy estimation, we observed substantial variation in chromosomal content across the species (**Figure 1e**, **Figure S4**). Strains from clades A and C were predominantly haploid and, when diploid, were highly homozygous. Only a few isolates in clade C exhibited low levels of heterozygosity, consistent with inbreeding. In contrast, clade B strains showed greater genome plasticity, including polyploidy, pronounced aneuploidies, and substantial heterozygosity. Strains from dairy fermentations exhibited especially high levels of heterozygosity, likely accumulated during prolonged asexual propagation under industrial conditions. The elevated ploidy and frequent aneuploidies observed in clade B are consistent with hallmarks of yeasts adapted to anthropogenic environments [14,18]. Together, these results define a species-wide structure of three major lineages only weakly associated with broad geographic origin, and identify clade A as a broadly distributed natural background against which clade B and clade C stand out for contrasting genomic signatures.

### An agave-restricted lineage shows deep genome-wide nucleotide divergence

The population structure analyses above identified clade C as a highly divergent lineage confined to agave fermentations. To determine whether this divergence reflects a genome-wide discontinuity, we compared clade C with the other *K. marxianus* lineages using *de novo* genome assemblies and estimating average nucleotide identity (ANI), gene-tree concordance, and synteny conservation. The exceptional divergence of clade C was already apparent in the first *K. marxianus* genomic phylogeny based on 17 strains, even though only a single isolate from this lineage was available at that time. Nevertheless, the ITS sequences of all clade C strains shared over 99% identity with the CBS 712T type strain (clade A), supporting their classification as *K. marxianus* isolates (**Table S2**).

*de novo* genome assemblies from clades A and C consisted of 46 to 200 contigs and showed 97 to 98% BUSCO completeness (**Figure S5**). Those from clade B were more fragmented and had lower scores, likely reflecting the greater difficulty of assembling heterozygous genomes with higher ploidy. Across the collection, the average genome size was 10.7 Mb and the mean number of annotated genes per genome was 4,900. These estimates are very similar to those of the chromosome-level DMKU3-1042 reference genome (10.9 Mb and 4,941 genes).

Whole-genome comparisons further highlighted the sharp separation of clade C. ANI between genomes from clades A and B exceeded 98% (**Figure 2a**), consistent with their relatively close relationship in the phylogeny and with the possible intraspecific hybrid origin of strains from clade B. In contrast, comparisons involving clade C yielded substantially lower ANI values, with averages of 95.6% between clades A and C and 95.2% between clades B and C (**Figure 2a**). These values fall just below the 95.7% threshold proposed as a species boundary for yeasts in the genus *Kluyveromyces* [39]. Comparison with the closest known sister species, *K. lactis* (NRRLY-1140), yielded much lower ANI values, close to 75%.

**Figure 2.**
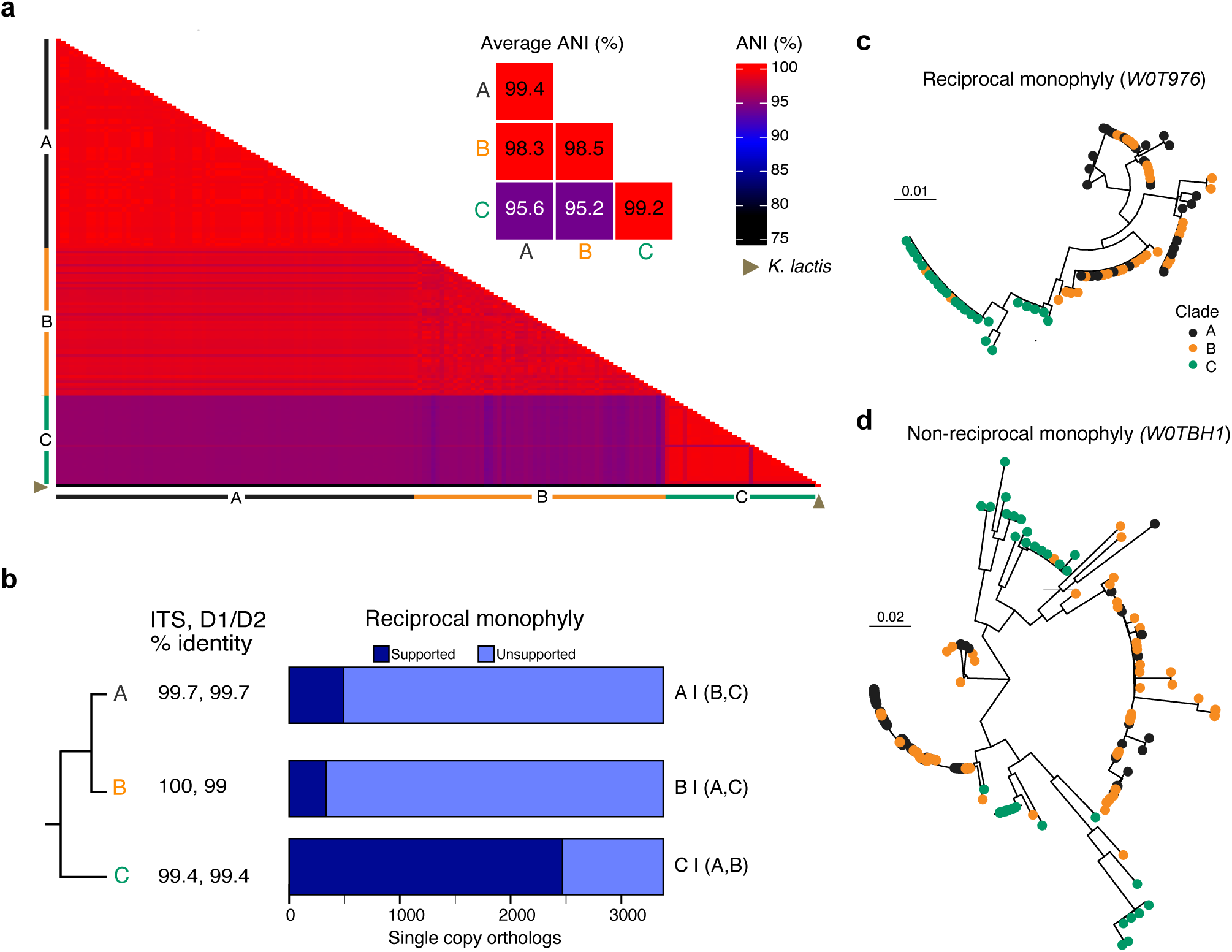
Genome-wide nucleotide divergence and widespread reciprocal monophyly define the agave-associated lineage. **a)** Heat map on the left displays ANI scores between all pairs of *K. marxianus* genomes as indicated by the color legend. Strains are sorted by the phylogenetic clade to which they belong (Figure 1b) as indicated by the color bars at both sides of the heatmap (A, black; B, orange; C, green). The genome of *K. lactis* was included as a reference at the bottom (brown triangle, ANI=75%). The heat map on the right shows the average ANI score between strains of the three clades. **b)** Mean nucleotide identity in the ITS and D1/D2 ribosomal regions of the three phylogenetic lineages relative to the CBS 712T type strain. Right, number of single-copy orthologs from the core genome (*n*=2,468) that support reciprocal monophyly of the three phylogenetic clades. **c,d)** Gene trees of the V-type proton ATPase subunit D (*W0T976*) and the tRNA (cytosine-5-)-methyltransferase Ncl1 (*W0TBH1*), illustrating the presence and absence of reciprocal monophyly, respectively

We next asked whether this deep divergence is also reflected in gene-tree concordance across the genome. Reciprocal monophyly among gene trees is an important criterion in species delimitation because it suggests long-term isolation and independent evolutionary trajectories among diverging lineages. To assess this, we quantified the number of OrthoFinder gene trees that support reciprocal monophyly of clade C (**Figure 2b**, **Figure S6**). We analyzed 3,377 single-copy orthologue gene trees using both genetic distance- and topology-based approaches (see Materials and Methods). Most gene family trees (2,468) consistently recovered clade C as a distinct monophyletic lineage, whereas the remaining trees were largely uninformative, likely due to incomplete lineage sorting (examples in **Figure 2c,d**). Comparisons of intra- and inter-clade genetic distances further indicate that, for nearly all genes, sequences in clade C are more divergent from those in clades A and B than from each other within clade C (**Figure S6**).

To test whether the divergence of clade C could instead reflect large structural rearrangements, we re-sequenced representative strains from each clade using long-read technology. We selected strains that were haploid from a shared fermentative environment, capturing genetic differences among clades. Genomes were assembled to near chromosome-level resolution (9-10 scaffolds) and mapped to the draft reference genome NBRC1777 (clade A). Over 95% of the orthologous genes from NBRC1777 (5,080 genes) were identified across the three genomes. A total of 4,862, 4,993, and 4,874 genes were annotated in YMX005664 (clade A), YMX009855 (clade B), and YMX005668 (clade C), respectively. Despite their nucleotide divergence, the genomes were highly collinear, with only minor structural differences (**Figure S7**). Synteny analysis revealed a similar number of translocations (∼17-20 events) and duplications (∼18-23 events) in the genomes of the three strains, most of them near telomeric regions. Only a few inversion events were detected, and these were shared among lineages, indicating that they predate their divergence. These results suggest that the genomic evolution of clade C is associated primarily with the accumulation of single-nucleotide changes rather than with major structural rearrangements.

Together, these analyses show that the agave-restricted clade C is separated from the rest of *K. marxianus* by deep genome-wide nucleotide divergence, yet retains broad structural collinearity. This combination defines clade C as a sharply differentiated lineage within the species.

### Reduced mating compatibility supports partial reproductive isolation in the agave-associated lineage

The comparative genomic analyses above identified clade C as a deeply divergent lineage within *K. marxianus* despite its coexistence with clade A in agave fermentations. To determine whether this genomic discontinuity is accompanied by reduced reproductive compatibility, we examined sexual reproduction, mating efficiency, and patterns of allele sharing among the three major lineages. Because strains from clades A and C are mostly haploid, but the sequenced set included diploid isolates from both lineages, we first assessed their ability to complete sexual development by measuring sporulation efficiency and spore viability (**Figure 3a**). Isolates from clade B generally show higher ploidy, but we included only diploid strains in these assays because aneuploidies could impair spore viability. Altogether, we analyzed four diploid isolates each from clades A and C, along with 14 from clade B (**Figure 3b,c**). Sporulation assays were performed under a variety of media, induction temperatures, and agitation regimes previously described for *K. marxianus* and related yeasts (**Table S3**). However, all strains that were able to sporulate did so most efficiently in 2% potassium acetate at 30 °C, with no clear clade-specific preference. Among the 22 diploid strains examined, only eight were capable of sporulation, and overall efficiencies were low: between 6 and 12% for clade A, between 2 and 18% for clade B, and fewer than 4% of cells formed tetrads for the single clade C strain that sporulated (**Figure 3c**).

**Figure 3.**
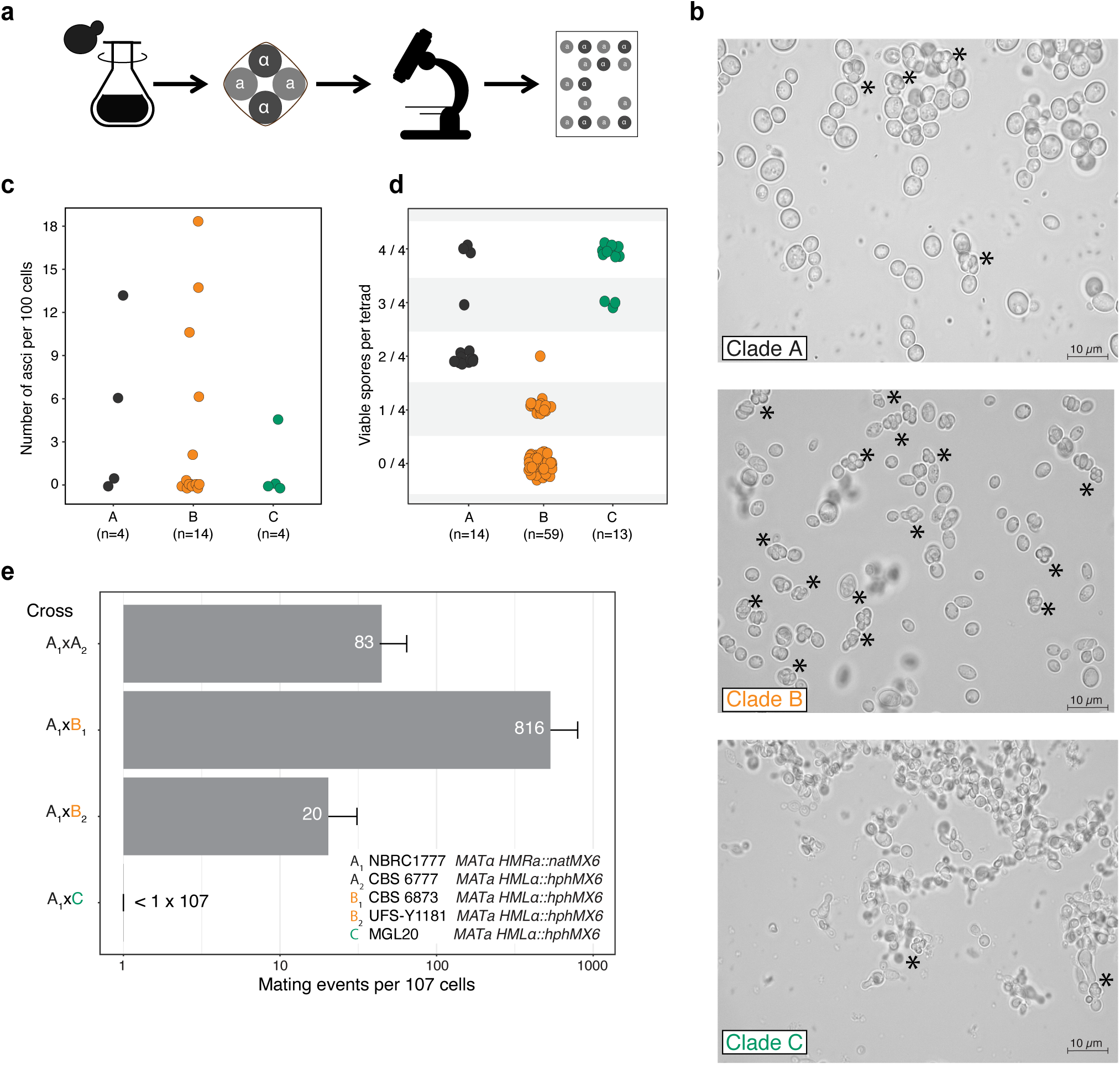
Sexual reproduction and mating compatibility differ among the major lineages of *K. marxianus*. **a)** Schematic of the characterization workflow for diploid isolates used in sporulation efficiency and spore viability analyses. **b)** Light microscopy images of sporulating representative diploid strains from clades A (94.204-1), B (CBS_1574), and C (YMX005681). Sporulation was induced in 2% potassium acetate at 30°C for two days with constant shaking. Asterisks indicate asci. **c)** Sporulation efficiency of the 22 available diploid strains from the three clades. Values represent the percentage of total cells that formed asci, counting at least 100 cells per visual field. Strains are grouped by clade, as indicated by the color of the dots (A, black; B, orange; C, green) and the number of strains analyzed for each clade is shown in parenthesis. **d)** Spore viability across the eight strains that were able to sporulate. A spore was considered viable if it formed a visible colony after tetrad dissection; only strains capable of forming asci with four spores were dissected. The number of viable spores per dissected tetrad is shown, with the total number of dissected tetrads for each clade indicated in parentheses. e) Inter- and intra-clade crosses used to assess mating compatibility between subpopulations. Values in the bar plot represent the number of colonies that grew on YPD supplemented with the two antibiotics, nourseothricin and hygromycin B, per 10 million cells used in each cross. *MAT*α parental strains were marked with the natMX6 nourseothricin resistance cassette, while *MATa* parental strains carried the hphMX6 hygromycin B resistance marker.

To assess spore viability, asci from all eight strains that sporulated were dissected under the microscope. Despite their higher sporulation capacity, strains from clade B showed the lowest spore viability, with one to zero surviving spores per tetrad. Strains from clade A yielded two to four viable spores, and all dissected asci from the single sporulating strain from clade C contained three to four viable spores (**Figure 3d**). The low spore viability of clade B is consistent with the possible intraspecific hybrid origin of this lineage and with the industrial nature of these isolates. Human management may have contributed to reduced fertility, as has been documented in other domesticated yeast populations. At the same time, these results indicate that isolates from all three clades retain at least some capacity for sexual reproduction.

We next evaluated mating efficiency among *K. marxianus* isolates. To do so, we replaced the silenced mating loci *HMRa* and *HML*α with the dominant drug-resistance markers *natMX6* and *hphMX6*, respectively. This strategy allowed us to detect rare mating events by selecting for growth on medium containing both nourseothricin and hygromycin B, while also preventing mating-type switching. After determining the mating type of haploid strains to identify compatible pairs, we found that the majority were MATα, regardless of clade. Because most clade B isolates were diploid, we used viable spores recovered from the sporulation assays. Despite the mating-type imbalance, and after several transformation attempts, we successfully obtained stable *MATa* strains from all clades, but were able to transform *MAT*α strains only from clade A. As a reference, we performed mating assays with strains of the sister species *K. lactis*, which yielded a mating frequency of approximately 1.0×10^-7^. This was about tenfold lower than previously reported for that species, but we note that the strains used here were different [40].

Using these marked strains in three independent mating assays, we observed substantial variation in mating frequencies within and between phylogenetic groups. Crosses between two clade A strains (NBRC1777 x CBS 6777) yielded, on average, a mating frequency of 8.3×10^-6^ (**Figure 3e**). Interclade crosses between clades A and B showed a range of outcomes: a cross between two natural strains (NBRC1777 x CBS 6873) exhibited the highest mating frequency, at approximately 8.0×10^-5^, whereas the cross between the natural strain NBRC1777 and the beer strain UFSY-1181 yielded a mating frequency below 2.0×10^-6^. In contrast, no detectable mating events were observed in any cross involving clade C strains, suggesting that mating frequencies in these combinations fall below the 1.0×10^-7^ detection limit. Thus, although clade C retains the ability to complete meiosis when present in a diploid background, its mating compatibility with the other major lineages appears strongly reduced.

To further assess the apparent isolation of clade C, we examined patterns of gene flow using Patterson’s *D* statistic, which quantifies asymmetries in shared ancestral polymorphisms through ABBA and BABA site counts across different subpopulation quartets. In all tested scenarios, the two clade C subpopulations (Pop10 and Pop11) showed *D* values consistent with minimal allele sharing with subpopulations from clades A and B (**Figure S8**, **Table S4**). The opposite pattern was observed among subpopulations within clades A and B, consistent with either recent gene flow or very close shared ancestry. Together, the marked reduction in detectable mating involving clade C and the limited allele sharing between this lineage and the rest of the species support partial reproductive isolation of the agave-associated lineage represented by clade C.

### A domesticated lineage shows genome remodeling, aneuploidy, and erosion of sexual reproduction genes

In contrast to the deep nucleotide divergence that characterizes clade C, the ploidy analyses described above identified clade B as the lineage with most extensive genome remodeling within *K. marxianus*. This clade is composed predominantly of strains from dairy and beer fermentations, together with a small number of clinical isolates, linking its distinctive genomic profile to recurrent occupation of human-managed environments. To define the genomic features underlying this domestication-associated trajectory, we examined gene-content variation and copy-number change across the three major lineages.

We first reconstructed the *K. marxianus* pangenome using protein annotations from strains of all clades with medium-to-high BUSCO completeness (Materials and Methods). In total, we identified 4,908 Orthologous Gene Clusters (OGCs) in the species pangenome. Of these, 3,873 genes constitute the core genome, shared by all strains, while the remaining 1,035 genes form the accessory fraction, exhibiting variable presence across strains (**Figure 4a**). Comparing *K. marxianus* OGCs with those of its sister species *K. lactis* and the model yeast *S. cerevisiae*, we found a high conservation rate, with 83.5% (4,292/5,136 OGCs) shared among the three species. Most of these conserved genes belong to the *K. marxianus* core genome, whereas most *K. marxianus*-specific genes are annotated as hypothetical proteins. Notable exceptions include homologs of genes with known functions that appear duplicated in this species, such as a homolog of *GAL4*, which encodes a transcriptional regulator of the galactose pathway, and homologs of *COX1*, a component of the mitochondrial electron transport chain. At the chromosome level, these losses were widely distributed across the genome in clade B rather than concentrated on one or a few chromosomes (**Figure 4b**). Clade C showed a weaker but still detectable signal of losses at multiple chromosomal positions, whereas clade A displayed only sparse events.

**Figure 4.**
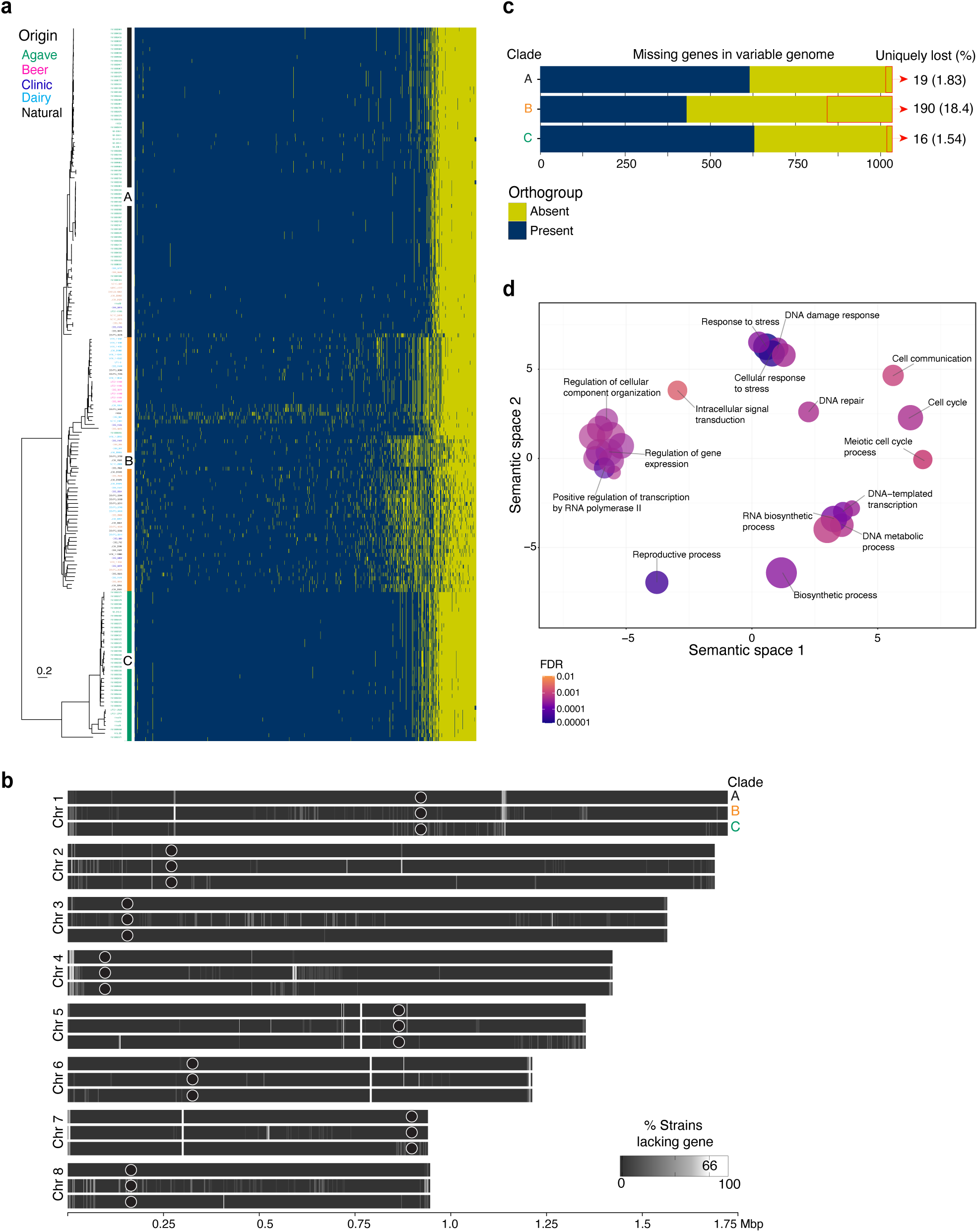
A domesticated *K. marxianus* lineage shows gene loss across the genome and erosion of sexual reproduction genes. **a)** Presence-absence matrix of orthologous gene clusters (OGCs) across strains from the three phylogenetic clades (A, black; B, orange; C, green). Rows represent genomes, columns represent OGCs in the variable genome (*n*=1,035), and colors indicate presence (blue) or absence (yellow). **b)** Chromosome-level distribution of genes lost in each clade. For each locus, the percentage of strains sharing the loss is shown as a color gradient; the lightest regions represent genes lost in at least two-thirds of the genomes within that clade. Black circles indicate the centromeres, as described by Ortiz-Merino et al. [30]. **c)** Number of genes from the variable genome (*n*=1,035) lost in each clade. Uniquely lost genes, defined as absent only in one clade and not in the other two, are indicated at the right of each bar with red rectangles. **d)** Gene Ontology (GO) enrichment analysis of Clade B-specific OGC losses. GO annotation was based on *S. cerevisiae* orthologs identified with OrthoFinder. Enriched categories are projected onto semantic spaces 1 and 2, whereby clustered circles represent functionally related categories. Circle size is proportional to the number of genes assigned to each GO category; only select categories are labeled.

We next asked whether gene-content changes are distributed evenly across the three lineages. To investigate lineage-specific gene erosion at a broad scale, we defined gene losses as OGCs absent in at least 75% of the strains within a given clade. Clade B exhibited a striking reduction in gene content, with 190 genes from the pangenome lost, compared with only 19 and 16 genes lost in clades A and C, respectively (**Figure 4c**). Thus, despite its high nucleotide divergence, clade C retains a largely conserved gene repertoire, whereas the broad gene loss observed in clade B is consistent with the more extensive genome remodeling that distinguishes this domesticated lineage. Functional analysis of the genes lost in clade B, including Gene Ontology (GO) enrichment, revealed a marked depletion of genes involved in sexual reproduction (**Figure 4d**, **Table S4**). For example, *IME2* and *IME4*, key regulators of pre-meiotic initiation, chromosome segregation, and progression through meiosis, were absent in a substantial fraction of dairy-associated strains from this clade. Other missing genes included *RIM4*, a positive regulator of sporulation-specific genes, and *SPO14*, essential for meiosis and spore formation. The absence of these genes is consistent with the poor sporulation capacity and low spore viability observed in clade B strains.

Additional gene losses in clade B affected regulators at multiple functional levels, including transcription (*CUP9*, *BAS1*, *DAL80*), chromatin remodeling (*PHO23*, *SDS3*, *DOT1*), and post-translational modification processes such as ubiquitination (*UBP5*, *APC11*, *LEE1*), glycosylation (*MNN14*, *MNT4*), and phosphorylation. Although these regulators span diverse biological processes, stress response was notably enriched among them. Losses also included enzymes involved in cofactor biosynthesis, as well as lipid and galactose metabolism. As with the genes related to sexual reproduction, these losses point to genome restructuring likely associated with adaptation to industrial conditions.

To further characterize genome remodeling in clade B, we analyzed copy number variation (CNV) across the three major phylogenetic clades using read-depth based copy-number inference with ploidy-normalized coverage [41] (**Figure S9**, **Table S6**). Clade B genomes exhibited a higher frequency of duplications, with 98 loci showing increased copy number not observed in the other clades. In addition, 35 loci in clade B displayed reduced coverage, consistent with the gene losses identified above. Clade C also showed 101 clade-specific CNVs, whereas no clade-specific CNVs were detected in clade A. However, in clade C these variants were not accompanied by the extensive gene loss, elevated ploidy, recurrent aneuploidy, or reduced fertility that distinguish clade B. Together, the extensive gene loss, CNV enrichment, aneuploidy, and reduced fertility of clade B define a domestication-associated trajectory of genome remodeling within *K. marxianus* that is distinct from the nucleotide-based divergence observed in the agave-associated clade C.

### Phenotypic divergence reflects lineage structure and environmental association

The observed genomic and reproductive contrasts led us to ask whether lineage divergence is also reflected at the phenotypic level, particularly in traits relevant to environmental performance and biotechnological use. To address this, we profiled growth variation across the sequenced *K. marxianus* collection under a broad set of environmental and nutritional conditions. We measured growth rates for all strains not protected by proprietary rights in the presence of different temperatures, carbon sources, chemical stressors, and salts. Clustering strains by their performance across conditions broadly recapitulated the three major phylogenetic clades, with strains from clades B and C grouping especially closely (**Figure 5a**).

**Figure 5.**
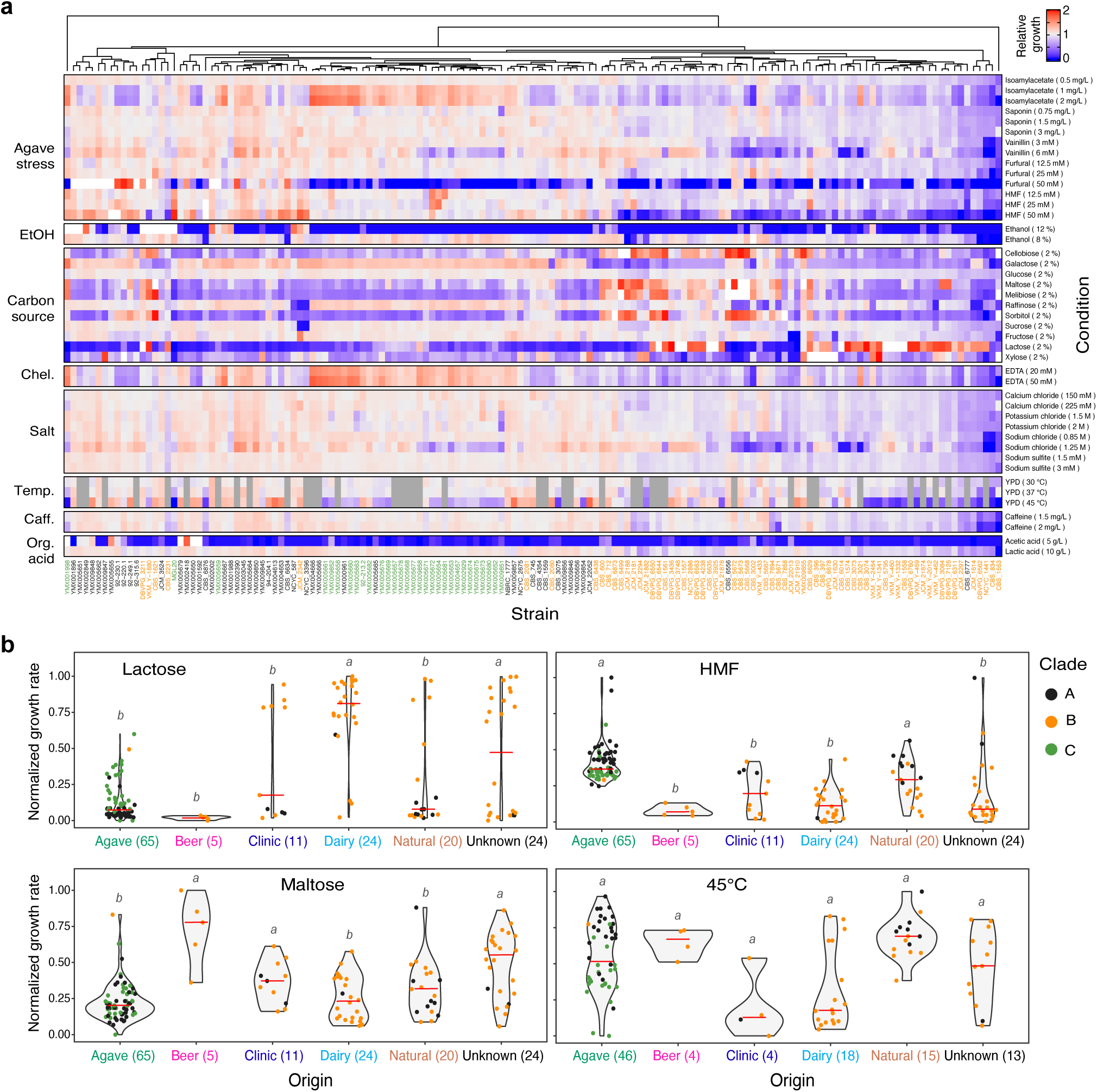
Phenotypic divergence reflects lineage structure and environmental association. **a)** Heat map of growth profiles for a subset of the sequenced *K. marxianus* strains (*n*=136) across multiple conditions. Values represent the growth rate of each strain relative to the reference strain NBRC1777 in each condition; missing values are shown in gray. Columns correspond to strains, grouped by hierarchical clustering based on the Euclidean distance of their phenotypic profiles using complete linkage. The color of each strain label indicates its phylogenetic clade (A, black; B, orange; C, green). Rows correspond to growth conditions, grouped by stress type. **b)** Growth rates in lactose (upper left), maltose (lower left), in the presence of HMF (upper right), and at 45 °C (lower right), grouped by strain origin and normalized to the maximum observed growth rate in each condition. The number in parathesis for each origin show the number of strains.

Despite this overall phylogenetic signal, the different lineages also showed distinct phenotypic biases consistent with their environmental associations. In general, strains from clade B grew more slowly in the presence of stress factors such as saponin, furans, and osmotic stressors, in agreement with their anthropogenic origin (**Figure 5b**). However, clade B strains showed superior growth under other conditions, most notably when lactose was provided as a carbon source. Dairy-associated isolates from this clade displayed the highest growth rate on lactose, reaching a maximum of 0.28 h^-1^, around ten-fold higher than strains from clades A and C and consistent with previous reports on dairy *K. marxianus* strains [21,42]. In addition, beer strains from clade B showed the highest capacity for maltose utilization, with growth rates of 0.15 h^-1^. This agrees with observations in beer populations of *S. cerevisiae* [43] and suggests convergent adaptation to brewing environments. Proficient growth at 45 °C was detected across all three clades, consistent with thermotolerance as a broadly conserved trait in *K. marxianus*. Such growth rates nevertheless varied substantially among strains, indicating quantitative variation in thermotolerance that was not significantly structured by environment of origin.

Importantly, strains from agave fermentations shared phenotypic traits linked to that environment despite belonging to different clades. In particular, agave isolates from clades A and C showed increased growth compared to other environments of origin in the presence of furans such as furfural and hydroxymethyl furfural (**Figure 5b**), which are oxidative stress-inducing compounds generated during agave cooking that inhibit microbial growth [44]. Thus, two otherwise distinct evolutionary backgrounds, the broadly distributed clade A and the deeply divergent clade C, converge on enhanced growth under a stress condition directly associated with agave fermentations. Phenotypic profiles therefore reflect both lineage history and environment-associated convergence.

Together, these results show that the lineage structure defined by genomic analyses has phenotypic consequences, with direct implications for the identification and selection of strains suited to different biotechnological applications. Overall, our results indicate that diversification in *K. marxianus* extends from genome structure to performance across environments, resulting in lineages that differ not only in ancestry but also in functional potential.

## Discussion

*K. marxianus* is a yeast of growing interest as a chassis for biotechnological applications, yet its intraspecific diversity has remained relatively underexplored. More broadly, it offers a useful system for addressing a central problem in population genomics, namely whether genetic diversity within a species is best interpreted as continuous variation, as discrete lineage separation, or as an intermediate state in which gene flow, ecological association, and reproductive compatibility are partly decoupled [6,8,45]. Our results argue against a simple continuum of diversity in *K. marxianus*, but also against a binary view in which all major lineages represent equivalent steps toward species separation. Instead, the species contains distinct lineage-level outcomes produced by different evolutionary processes. Clade B illustrates a domestication-associated trajectory marked by genome remodeling, altered reproductive performance, and limited sequence divergence from clade A. By contrast, clade C represents a deeply divergent, agave-associated lineage with little detectable recent gene flow and reduced reproductive compatibility. Thus, *K. marxianus* shows how marked genomic discontinuities can emerge within a single species through more than one evolutionary route.

This conclusion is strengthened by the weak association between lineage structure and broad geographic origin. Strains from different lineages often occur in the same regions and, in some cases, in the same fermentative environment. This was especially evident in agave fermentations, which occur across a broad region of Mexico and harbor most of the detected subpopulation diversity. This pattern contrasts with other yeasts in which broad biogeographic structure is more clearly reflected in genome-wide diversity [12]. *K. marxianus* therefore provides a useful case in which lineage discontinuities cannot be explained simply as broad geographic subdivision, but instead reflect a combination of domestication-associated genome change, deep lineage divergence, restricted gene flow, and differences in reproductive compatibility.

Our sequencing effort resolves the status of the highly divergent agave-associated lineage previously represented by a single isolate from South Africa [30]. Rather than representing the extreme end of an undersampled continuum, clade C emerges here as a sharply differentiated lineage that remains exclusive to agave fermentations despite extensive new sampling. Clade C forms a long, separate branch in the phylogeny, contributes a disproportionate fraction of the species-wide SNP diversity, shows ANI values approaching proposed species-delimitation thresholds in *Kluyveromyces*, and is recovered as reciprocally monophyletic across much of its genes. At the same time, long-read comparisons indicate that this divergence is not driven by large-scale structural reorganization, but primarily by the accumulation of nucleotide-level differences across an otherwise collinear genome. Together, these features suggest that clade C is unlikely to represent a recently derived, locally specialized population and instead support its interpretation as a deeply divergent lineage within the species.

An additional striking feature of clade C is that all known isolates come from traditional agave fermentations. These environments are already known to harbor substantial yeast diversity at both the species and population levels [35,37,46,47]. Clade C is therefore unusual not simply because it comes from a diverse fermentation system, but because it is both the earliest-diverging lineage detected in *K. marxianus* and the lineage most narrowly associated with that environment. This pattern does not reveal whether agave fermentations represent the long-term environment in which this early-diverging lineage arose. Given the long biocultural history of agave use in Mesoamerica [48,49], where agaves originated and diversified, a long-term association is plausible. Alternatively, agave fermentations may have preserved a relict branch of *K. marxianus* that is now absent from, or still undetected in, other environments. In either case, clade C coexists locally with clade A, and more occasionally with clade B, yet shows little evidence of admixture or recent gene flow with the rest of the species, consistent with partial reproductive isolation despite sympatry.

In contrast, clade B shows a different evolutionary trajectory. This lineage is dominated by strains from recurrent human-managed environments, especially dairy and beer fermentations, whereas clinical isolates constitute a minor fraction in the linages and do not alter this overall ecological pattern. Although it remains relatively close to clade A at the nucleotide level, it shows extensive genome remodeling, including increased ploidy, frequent aneuploidies, abundant CNVs, and pervasive gene loss. These features are most pronounced in dairy- and beer-associated strains and are consistent with genomic patterns often linked to yeast domestication [12,14,18]. Yet unlike clade C, clade B does not appear fully reproductively isolated from the rest of the species, as at least some strains retain the capacity to mate with clade A. Clade B therefore exemplifies a domestication-associated trajectory in which genome architecture and reproductive performance have been altered without comparable deep nucleotide divergence.

Whereas *S. cerevisiae* shows extensive admixture among wild and domesticated lineages, including lineages with introgressed ancestry [12], gene exchange in *K. marxianus* appears more restricted, occurring mainly between the closely related clades A and B, while clade C remains largely separated from the rest of the species. This difference may partly reflect the predominantly haplontic life-cycle described for *Kluyveromyces*, in which starvation promotes mating-type switching, mating, and subsequent sporulation [30,50,51]. In contrast, *S. cerevisiae* has a primarily diplontic life-cycle, where diploid cells proliferate mitotically and enter meiosis under starvation. In this context, reduced mating between clade C and the other lineages, together with poor sporulation and spore viability in clade B, may further limit gene exchange in *K. marxianus*.

The phenotypic data are broadly consistent with this dual view of *K. marxianus* diversification. Growth profiles recapitulate the major phylogenetic groupings, indicating that lineage history is reflected at the phenotypic level. However, strains from shared environments also show convergent traits that likely reflect repeated exposure to similar selective pressures. Dairy- and beer-associated isolates from clade B perform especially well on lactose and maltose, respectively, consistent with adaptation patterns described in domesticated *Saccharomyces* lineages [52]. Likewise, agave-associated strains from clades A and C show increased tolerance to furans, compounds generated during agave cooking that inhibit microbial growth [44]. Clade A nevertheless retains broader stress tolerance, including stronger thermotolerance, and thus appears to preserve a more generalist phenotype than the more specialized industrial lineage.

These patterns also have practical implications for strain selection, the interpretation of genotype-phenotype relationships, and the design of crossing or engineering strategies in biotechnology. Recent comparative work has shown that *K. marxianus* stands out within *Kluyveromyces* for tolerance to high temperature and several chemical stresses [53]. Moreover, a recent study to improve lactic-acid production showed that screening broad strain diversity, including isolates from underrepresented environments, can uncover superior engineering backgrounds [28]. Notably, when viewed in the context of the population framework developed here, the strongest candidates for lactic-acid production fall within the agave-associated clade A rather than within the domesticated clade B. This indicates that industrially valuable traits are not necessarily concentrated in the most employed domesticated strains.

Together, these findings position *K. marxianus* as a useful system for studying how intraspecific diversity can extend from continuous variation to discrete lineage separation. In this species, some lineages remain connected by limited admixture despite domestication-associated genome change, whereas others are separated by sharper genomic discontinuities and reduced reproductive compatibility. By integrating population genomics, reproductive assays, and phenotypic profiling, our study provides a framework for studying intraspecific diversification and for exploiting the lineage-specific biotechnological potential of *K. marxianus*.

## Materials and Methods

### Strains, media, and DNA extraction

The *K. marxianus* strain set included 86 strains from the YMX-1.0 culture collection [35], five previously described by [54], all associated with agave fermentations in Mexico, and 75 strains from culture collections (WI-KNAW Culture Collection-CBS, Industrial Yeasts Collection-DBVPG, National Collection of Yeast Cultures-NCYC, Japan Collection of Microorganisms-JCM, NITE Biological Resource Center-NBRC, and Russian Collection of Microorganisms-VKM). The culture-collection strains include isolates from dairy products (cheese, yoghurt), beer, decaying foods, and clinical sources (mastitis, tuberculosis), as well as soil, water, and diverse environmental samples. The metadata for the strains used are provided in **Table S1**. DNA extraction was carried out from saturated cultures in YPD broth (10 g/L yeast extract, 20 g/L bactopeptone, 20 g/L glucose) grown at 30°C. Genomic DNA was purified with the MasterPureTM Yeast DNA Purification Kit, according to the manufacturer’s instructions. DNA samples were quantified by Qubit^TM^, and quality was assessed with a NanoDrop spectrophotometer.

### Short-read genome sequencing, assembly, and annotation

Libraries were prepared with the MGIEasy FS DNA Library Prep kit and sequenced on the DNBSEQ platform by the Beijing Genomics Institute-BGI (Guangdong, China). Base calling was performed using *Zebracall* (BGI), which converted raw signal data into FASTQ files [55,56]. For quality control, sequencing adapters were trimmed and low-quality reads were removed with *fastp v0.20.0*, generating clean reads for *de novo* assembly [57]. Genomes were assembled using *SPAdes v3.12.0* [58], and assembly quality metrics were evaluated with *QUAST v5.2.0* [59]. The final assemblies were used for genome annotation. For structural annotation, transcript and protein files from the public chromosome-level assembly of *K. marxianus* strain DMKU3-1042 [60] were used to run *MAKER v3.01.3*. To refine annotations, additional rounds were performed with the *ab initio* gene model predictor SNAP (Semi-HMM-based Nucleic Acid Parser) [61]. *BUSCO v5.6.5* was run to assess genome completeness and generate retraining parameters for *AUGUSTUS v3.3.2*, starting with available gene models for *K. lactis* in the saccharomycetes_odb10 dataset [62]. Finally, *BLAST+ v2.7.1* was run to incorporate functional annotation for predicted proteins using homology with the UniProt database [63].

Ten previously available genomic datasets from NCBI were also included for downstream comparative genomic analysis. These comprised five genomes from [30], one genomic library from the diploid agave-fermentation strain SLP1 [33], and four genomes from strains isolated during henequen spirit production [31] (**Table S1**).

### Oxford Nanopore sequencing and assembly

A representative haploid strain from each phylogenetic clade was selected for long-read sequencing based on BUSCO completeness (> 98%) in the corresponding short-read assembly. The genomes of strains YMX005664 (clade A), YMX009855 (clade B), and YMX005668 (clade C) were sequenced using Oxford Nanopore Technologies on a MinION flow cell. Raw reads were trimmed and filtered with *Filtlong v0.2.0* using default parameters except for *--target_bases* 440000000, to retain reads corresponding to approximately 40-fold genome coverage, and *--min_length* 1000 [64]. For *de novo* assembly, *Canu v2.2* was used with default settings, setting the genome size to 11m [65]. Resulting assemblies were polished with the corresponding short-read libraries using *Racon v1.4.3* [66]. The resulting contigs were evaluated with *QUAST v5.2.0*.

### Variant calling, phylogenetic inference, and *in silico* ploidy estimation

A total of 178 genomic libraries (newly generated genomes and 10 public libraries) were used for variant calling. First, trimmed reads were mapped with *BWA v0.7.4* [67] to the haploid reference genome NBRC1777v1.0, selected for its high-quality assembly and low SNP density [30,68]. Unaligned reads were removed, and PCR-duplicated reads were marked with *Picard v2.6.0* (https://broadinstitute.github.io/picard). Variant calling was performed across all genomes with *GATK v4.1.1.0* HaplotypeCaller [69], following the Broad Institute best-practice workflow (https://software.broadinstitute.org/gatk/best-practices). Raw variants were first restricted to SNPs using *SelectVariants* and then filtered y *VariantFiltration* using the following criteria: Quality by Depth QD < 2.0, Mapping Quality MQ < 40, Fisher Strand Bias FS > 60, Strand Odds Ratio SOR > 3, Mapping Quality Rank Sum MQRankSum > −12.5, and Read Position Rank Sum > −8.0). *SelectVariants* was then used to retain only biallelic SNPs and allow a maximum of 10% missing data.

For phylogenetic reconstruction, a raw variant matrix of 1,345,626 variant sites was generated from the 178 genomic libraries. After filtering, a reduced matrix containing 953,880 biallelic SNPs was obtained and used for tree construction with *IQ-TREE v2.3.6* [70].

### Population genetic summary statistics

Genetic diversity (π) and Tajima’s *D* were estimated using *VCFtools v0.1.14* [71] on a VCF containing the variants that mapped to the *K. marxianus* NBRC1777 reference genome. The dataset was filtered to retain high-quality biallelic SNPs with less than 10% missing data. Per-variant heterozygosity was calculated using BCFtools v1.9 on individual strain VCFs by dividing, for each variant, the number of heterozygous genotype calls by the total number of genotype calls. Genome-wide *Fst* values were estimated for the different subpopulation clusters using the *SNPRelate v1.32.2* R package from Bioconductor [72].

### Population structure and ancestry inference

The *Admixture v1.3.0* model-based clustering algorithm was used to detect mixed ancestry and estimate the number of ancestral populations in the 178 sequenced genomes [73]. A set of 953,880 biallelic SNPs was pruned by removing markers in linkage disequilibrium (LD) with *PLINK v1.9*, using a 500-SNP window, a 10-SNP step size, and an *r*² threshold of 0.15 [74]. Admixture was run assuming between 1 and 30 ancestral populations (K = 1–30). Cross-validation errors generated by Admixture were used to determine the optimal subdivision of populations explaining genetic variation across the dataset. Population ancestry was visualized using *pong* [75] and the R package *pophelper v2.3.1* [76]. Principal component analysis (PCA) was performed with *PLINK v1.9*.

### Whole-genome average nucleotide identity

ANI scores between whole-genome *de novo* assemblies were calculated with *FastANI v1.34* [77]. Because shorter k-mer sizes are more sensitive for divergent genomes, ANI was calculated using a k-mer size of 22, as previously recommended for comparisons among divergent yeast genomes [39]. A total of 178 assemblies, including the chromosome-level *K. lactis* NRRL Y-1140 reference genome (GCA_000002515.2) as an outgroup, were analyzed to generate a matrix of 15,931 pairwise comparisons containing ANI scores as percentages of nucleotide identity. For visualization, ANI scores were sorted by phylogenetic clade.

### Reciprocal monophyly across orthologous gene trees

To assess the degree of reciprocal monophyly of clade C across orthologous gene trees, both topological and genetic distance-based approaches were applied using gene trees inferred with *OrthoFinder v3.0.1* [78]. Orthogroups were identified across all genomes through all-versus-all sequence similarity searches, followed by multiple sequence alignment and gene tree reconstruction using *FastTree v2.1.11*, yielding 4,908 gene trees. For the topological approach, a relaxed monophyly criterion was applied. Gene trees were parsed in R using the *ape* package *v5.7-1* to identify clade C tips and inspect internal nodes for clustering. A tree was considered to support relaxed monophyly if ≥90% of clade C sequences grouped within a single subclade, allowing minor deviations attributable to incomplete lineage sorting or reconstruction errors. In parallel, genetic cohesion was evaluated by computing pairwise intra- and inter-clade distances for all gene trees using *ape*. Distances were categorized based on clade comparisons (e.g., within-clade C vs. between-clade), and one-sided Wilcoxon rank-sum tests were applied to test whether within-clade C distances were significantly lower than between-clade distances. Resulting *p*-values were adjusted for multiple testing using the false discovery rate (FDR) method.

### Synteny and structural variant analysis

Genome-wide synteny between the reference *K. marxianus* NBRC1777 genome and the Oxford Nanopore–sequenced genomes was assessed using *MUMmer v4.1.1* [79] excluding mitochondrial sequences from the reference assembly. One-to-one alignments were retained using *delta-filter*, and alignment coordinates for all genes were extracted. Based on these alignments, contigs from each *de novo* assembly were assigned to reference chromosomes, oriented, and reordered according to their best syntenic matches. Gene-level synteny was then inferred by aligning reference CDS sequences against the reordered contigs of each query genome using *BLASTn*. *BLAST* results were processed in R to generate gene order and synteny anchor files, which were subsequently converted into *BED* and anchor formats compatible with the Python-based *MCScan* program. Final macrosynteny visualizations were generated using the *JCVI/MCScan* toolkit [80], producing chromosome-level karyotype plots that highlight conserved collinearity and structural variants, including inversions, duplications, and translocations, between each query genome and the reference.

### Gene flow inference using Pattersońs *D* statistic

To quantify allele-sharing asymmetries and assess evidence of historical gene flow among *K. marxianus* populations, Patterson’s D statistic was computed using the ABBA-BABA framework. A VCF containing genotypes for the 178 *K. marxianus* strains and the reference *K. lactis* CBS 683T strain, comprising 1,036,728 SNPs, was converted into *EIGENSTRAT* input format (*map, geno*, and *ind* files) using the script *convertVCFtoEigenstrat.sh* (rec = 2 cM/Mb) [81]. *D*-statistical analyses were performed with *ADMIXTOOLS v7.0.2* through the R package *admixr v0.9.1*, using the function *d* [82]. For each population quartet, D values were obtained together with standard errors, *Z*-scores, and counts of ABBA and BABA informative sites. Statistical significance was evaluated using block jackknife resampling, with an absolute *Z*-score ≥ 3 considered strong evidence for gene flow or excess ancestral allele sharing.

### Pangenome analysis and clade-specific gene loss inference

To characterize the species pangenome, orthologous gene clusters (OGCs) were inferred based on protein-coding gene predictions from all sequenced genomes. Protein FASTA files for each genome were used as input for *OrthoFinder v3.0.1*, which identifies OGCs by performing all-versus-all sequence similarity searches using BLAST [78]. The resulting orthogroups represent sets of homologous genes identified across the genome set. Based on these OGCs, genes were classified into two categories: core genes, defined as those present in all genomes, and accessory genes, defined as those exhibiting presence/absence variation across *K. marxianus* populations. To compare gene-content variation among clades, the proportion of genes considered lost within each clade was estimated by identifying genes absent in at least two-thirds of the strains from that clade, following a criterion previously used to detect pervasive gene loss in budding yeasts [83]. From the resulting gene-loss profiles, clade-specific gene losses were determined by excluding genes also absent from the other two clades.

### Gene ontology enrichment of clade-specific gene losses

To assess whether pervasive gene losses observed in each lineage were randomly distributed or enriched for functionally related gene groups, clade-specific gene-loss lists generated during pangenome reconstruction were used for GO enrichment analysis. Functional annotation was based on orthologs from *S. cerevisiae* (org.Sc.sgd.db), and statistical significance was assessed using Fisher’s exact test (*p* < 0.001) implemented in the R package *topGO v2.62* [84].

Hierarchical relationships among enriched Biological Process terms were visualized with *rrvgo v4.5* [85], which reduces semantic redundancy and represents GO terms in a scatter plot based on the first two principal coordinates of the semantic dissimilarity matrix.

### Read-depth-based copy-number variation inference

CNVs were identified by extracting read-depth information from BAM files, followed by signal aggregation, GC-content bias correction, and ploidy normalization. These steps were performed using *Control-FREEC v11.6* [41] with genomic window sizes of 500 bp and 1 kb. CNV regions were intersected with the annotated *K. marxianus* NBRC1777 reference genome, to which all reads had been previously mapped. Gene annotations were retrieved from the corresponding *GFF* file for the NBRC1777 reference genome using *GRanges* functions in the Bioconductor package *genomation v1.3.0*. For each gene overlapping a CNV region, copy number was extracted and classified as a duplication (>2 copies), deletion (zero copies), or neutral (single copy).

### Ploidy assessment by flow cytometry

Ploidy was assessed by flow cytometry following the Rosebrock method [86]. Briefly, a single colony was streaked onto YPD agar and incubated at 30 °C for 48 h. One colony was then inoculated into 5 mL YPD broth and grown overnight at 30 °C with shaking at 180 rpm. Cell density was determined from OD_600_, assuming that 1.0 OD_600_ corresponds to approximately 1.5×10^7^ cells/mL and verified using a Neubauer chamber. The starter culture was diluted to OD_600_ = 0.2 in fresh YPD and incubated at 30 °C and 180 rpm to mid-exponential phase (OD_600_ = 0.5–0.6). Approximately 1×10^7^ cells were harvested by centrifugation at 10,000 rpm for 5 min at 4 °C, the supernatant was removed, and the pellet was resuspended in 200 µL of 70% (v/v) ethanol by vortexing. Fixed cells were stored overnight at 4 °C or long-term at −20 °C.

For staining, fixed cells were thawed on ice, vortexed, and a 30 µL aliquot (∼1.5×10^6^ cells) was transferred to 270 µL of 70% ethanol. Cells were pelleted at 10,000 rpm for 20 min at 4 °C, washed twice with 50 mM sodium citrate buffer (pH 7.2), and resuspended in 300 µL of sodium citrate buffer containing RNase A (20 µg/mL) and SYTOX Green (1 µM; Invitrogen S7020). Samples were incubated for at least 1 h at 37 °C in the dark. Proteinase K (6 µL of 20 mg/mL) was then added, followed by incubation for at least 1 h at 55 °C or overnight at 4 °C. Stained cells were analyzed by flow cytometry at the lowest flow rate (12.5 µL/min), acquiring forward-and side-scatter area and width, and SYTOX Green fluorescence (FITC/GFP/YFP channel), all on a linear scale. FACS data were analyzed with *FCS Express^TM^ 7* software (De Novo Software, 2025).

### Determination of sporulation efficiency and spore viability

Sporulation assays were performed with diploid strains in four media: Minimal Sporulation medium (Spo) [87], Super Sporulation medium (Super-Spo), Uracil-Leucine sporulation medium (Spo-UL) [88], and YPA supplemented with potassium acetate (YPA+KOAc) [89]. Medium compositions are provided in **Table S3**. Cells were first grown in 3 mL YPD at 30 °C for 12 h. For Spo, Super-Spo, and Spo-UL, cells were harvested directly from YPD, washed twice with sterile water, and resuspended in the corresponding sporulation medium without pre-sporulation. For YPA+KOAc, cells underwent pre-sporulation in YPA medium containing potassium acetate: cells were transferred to 50 mL YPA at OD_600_ = 0.1, incubated at 30 °C for 6–8 h to OD_600_ ≈ 0.5 (∼10^7^ cells/mL), harvested, washed twice with sterile water, and resuspended in 50 mL of 2% potassium acetate to induce sporulation. All cultures were incubated at 30 °C with shaking at 300 rpm. Sporulation progression was monitored by light microscopy for up to 5 days; tetrads typically appeared after 24-36 h. Sporulation efficiency was quantified at 36 h by counting visible tetrads per 100 cells across multiple optical fields.

Spore viability was assessed by tetrad dissection of asci harvested 24 h after sporulation induction, a time point chosen to preserve tetrad integrity and minimize self-mating. A 10 µL aliquot of sporulated culture was mixed with 20 µL zymolyase solution (0.5 mg/mL) and spotted onto one end of a YPD agar plate. Individual asci were identified and dissected using an ECLIPSE Ci series microscope (Nikon) equipped with a micromanipulator. In each replicate, 18 tetrads per plate were dissected. Spore viability was determined from the ability of each spore to form a visible colony after incubation at 30 °C for 3 days.

### Mating-type verification and mating assays

Mating type was verified by colony PCR. Fresh cultures from frozen stocks were grown on YPD solid medium at 30 °C for 48 h, and gDNA was extracted by alkaline NaOH boiling preparation. PCR was performed using Phusion^TM^ DNA polymerase (ThermoFisher Scientific). The *MATa* locus (1062 bp) was amplified with primers YY270F-Fw (5’-TGCAACCAACCAATCCCTTCCAAATTC-3’) and YY271F-Rv (5’-TCTTCCTTGAACCCGAAGCAAAAGATC-3’), while the *MAT*α locus (1532 bp) was amplified with primers YY270F-Fw and YY272F-Rv (5’-AACTTCAATCCCCGACCCACCGCAGTC-3’). These primers were designed to amplify the transcriptionally active *MAT* locus on chromosome I, which defines mating type in *K. marxianus* [89].

Haploid strains unable to switch mating type were constructed for each clade by deletion of the *HMRa* and *HML*α loci for *MATa* and *MAT*α strains, respectively. These loci were replaced with the *natMX6* and *hphMX6* dominant drug resistance markers, respectively. Strain-specific deletion cassettes were generated by fusion PCR [90], containing the resistance marker with 350-400 bp flanking regions homologous to the 5’ and 3’ regions flanking *HMRa* and *HML*α. The cassette was concentrated with the QIAGEN MinElute PCR Purification Kit, and ∼1 µg was transformed by electroporation [91]. Transformants were plated on YPD with the corresponding drug, and colony PCR was used to confirm mutants by integration of the marker at the target locus. This strategy successfully transformed strains from various genetic backgrounds.

The mating-type-stable haploid strains were then used for mating assays. To this end, each parental strain was grown in YPD broth at 30 °C until reaching a density of 10^8^ cells/mL. Crosses were performed on solid malt extract medium (5 g/L malt, 30 g/L agar) for 3 days at 30 °C by placing 10^7^ cells of each parental strain on a cellulose filter, to facilitate recovery of most cells and enable quantification of mating frequency. Filters were then placed in 2 mL sterile tubes, cells were resuspended in water, and serial dilutions were used to quantify colony-forming units (CFU) on YPD plates supplemented with nourseothricin (100 µg/mL) and hygromycin B (200 µg/mL). Only diploid cells resulting from mating can grow on plates containing both drugs [92]. As a reference, mating frequency was estimated for the sister species *K. lactis*, using strains 155 (*MAT*α, *ade2*, *his3*, *ura3*) and 12/8 (*MATa*, *lysA*, *argA*, *ura3*) [93]. Mating assays were performed as described for *K. marxianus* strains, but successful mating was estimated by plating on SD medium without lysine and histidine, as the parental strains are auxotrophic mutants.

### Growth-based phenotypic profiling across carbon sources and stress conditions

Growth rates were estimated for 136 strains, excluding those subject to commercial or third-party restrictions on public phenotypic data release. Kinetic assays were conducted in 96-well plates using an automated robotic station (Tecan Freedom EVO200) integrated with a microplate reader (Tecan Infinite M1000), with optical density monitored every hour for 48 h. Growth in the presence of inhibitors, including ethanol, organic acids, EDTA, osmotic stressors, and stressors typical of agave fermentations environments [94] was estimated in YPD medium inoculated with precultures grown in the same medium to ∼2×10^7^ cells/mL. To test carbon source utilization, saturated cultures in complete YNB Glc (yeast nitrogen base 0.7%, glucose 2%) were used to inoculate plates containing YNB with different carbon sources at 2% concentration: xylose, glucose, fructose, lactose, maltose, melibiose, sucrose, raffinose, and sorbitol. All tested growth conditions are detailed in **Table S7**. Growth rate data for the 136 strains were automatically acquired during logarithmic phase [95] and expressed as growth rates relative to the reference strain NBRC1777. To compare multiple traits, relative growth rates were normalized by min-max scaling, ensuring that each condition was on the same scale and directly comparable between strains. Growth diversity was visualized with *ComplexHeatmap v2.14.0* [96] and *ggstatsplot v0.12.0* was used to compare phenotypic variation across strain origin [97]. Temperature tolerance was tested by measuring growth on YPD medium at three temperatures, 30, 37, and 45 °C, using a Bioscreen system (Bioscreen C, Labsystems, Helsinki, Finland). For this, 10 µL aliquots of YPD precultures were used to inoculate 200-well microtiter plates. Growth kinetics were recorded at OD_600_ every 30 min for 48 h. Relative growth was calculated relative to the growth rate at 30 °C. For all assays, control strains (CBS 6556, NBRC1777, CBS 397) were analyzed in triplicate, including the *S. cerevisiae* haploid strain BY4741 from the S288c background.

## Data availability

Genome sequencing data generated in this study have been deposited in the NCBI SRA under the BioProject accession PRJNA1419316. The accession numbers of each genome employed in this study, including those previously sequenced, are provided in Table S1.

## Author contributions

Conceptualization: KV, AD, EM; Methodology: JMM-H, JS, LFG-O, JAA-R, RAO-M, MRK, ALT-M, AM, LM; Data curation: JMM-H, JS, LFG-O, JAA-R, AM; Investigation: JMM-H, JS, LFG-O, MRK; Formal analysis: JMM-H; Visualization: JMM-H; Funding acquisition: LM, KV, AD, EM; Supervision: KV, AD, EM; Writing original draft: JMM-H, AD, EM; Writing, review and editing: all authors. All authors of this study have fulfilled the criteria for authorship, as their participation was essential for the design and implementation of the study.

## Acknowledgements

We thank Luis Aguilar and Santiago García for support and administration of computing resources, and Porfirio Gallegos Casillas, Susana Ruiz-Castro, Carina Uribe, Nancy Quintana-Rodríguez and Ivan Sedeño for technical assistance, Quinten Deparis for helpful discussions, and Roberto Coria, Anne Gschaedler, Marc-André Lachance, Carolina Henritta Pohl, and Joseph Schacherer for sharing strains and resources. We are grateful to all producers of agave spirits and fermented beverages who kindly contributed to this work by providing access to fermentation samples; acknowledgments for assistance with fieldwork and sampling by the YeastGenomesMx consortium are detailed in Gallegos-Casillas work [35].

## Funding

This work was funded by Secretaría de Ciencia, Humanidades, Tecnología e Innovación de México (Secihti) grants CF-2023-G-695 (LM, EM), CBF-2025-G-838 (LM, ADL, EM), PAPIIT IN212524 (LM), FWO cSBO projects Fucatil (HBC.2020.2623, KV) and Laplace (S004624N, KV), and the iBOF/21/092 POSSIBL funding (KV). EM was funded by Secihti for a sabbatical stay (I0200/111/2024).

## Competing interest

The authors declare no competing interests.

## Manuscript versions

This manuscript was first released as a pre-print at *bioRxiv* (Moreno-Hernández J. M., et al., 2026).

## Notes

### Competing Interest Statement

The authors have declared no competing interest.

## References

1. Sukumaran J, Knowles LL. Multispecies coalescent delimits structure, not species. Proceedings of the National Academy of Sciences. 2017;114: 1607–1612. doi:10.1073/pnas.1607921114

2. Campillo LC, Barley AJ, Thomson RC. Model-Based Species Delimitation: Are Coalescent Species Reproductively Isolated? Systematic Biology. 2020;69: 708–721. doi:10.1093/sysbio/syz072

3. Cadena CD, Zapata F. The genomic revolution and species delimitation in birds (and other organisms): Why phenotypes should not be overlooked. Ornithology. 2021;138: ukaa069. doi:10.1093/ornithology/ukaa069

4. Hausdorf B. Species Delimitation Using Genomic Data: Options and Limitations. Molecular Ecology. 2025;34: e17717. doi:10.1111/mec.17717

5. Singhal S, Leaché AD, Fujita MK, Cadena CD, Zapata F. A Genomic Perspective on Species Delimitation. Annual Review of Ecology, Evolution, and Systematics. 2025. 10.1146/annurev-ecolsys-102723-055311

6. Roux C, Fraïsse C, Romiguier J, Anciaux Y, Galtier N, Bierne N. Shedding Light on the Grey Zone of Speciation along a Continuum of Genomic Divergence. PLOS Biology. 2016;14: e2000234. doi:10.1371/journal.pbio.2000234

7. Ravinet M, Faria R, Butlin RK, Galindo J, Bierne N, Rafajlović M, et al. Interpreting the genomic landscape of speciation: a road map for finding barriers to gene flow. Journal of Evolutionary Biology. 2017;30: 1450–1477. doi:10.1111/jeb.13047

8. Stankowski S, Ravinet M. Defining the speciation continuum. Evolution. 2021;75: 1256–1273. doi:10.1111/evo.14215

9. Chambers EA, Lara-Tufiño JD, Campillo-García G, Cisneros-Bernal AY, Dudek DJ, León-Règagnon V, et al. Distinguishing species boundaries from geographic variation. Proceedings of the National Academy of Sciences. 2025;122: e2423688122. doi:10.1073/pnas.2423688122

10. Liti G, Carter DM, Moses AM, Warringer J, Parts L, James SA, et al. Population genomics of domestic and wild yeasts. Nature. 2009;458: 337–341. doi:10.1038/nature07743

11. Liti G. The natural history of model organisms: The fascinating and secret wild life of the budding yeast S. cerevisiae. eLife. 2015;4: e05835. doi:10.7554/eLife.05835

12. Peter J, De Chiara M, Friedrich A, Yue J-X, Pflieger D, Bergström A, et al. Genome evolution across 1,011 Saccharomyces cerevisiae isolates. Nature. 2018;556: 339–344. doi:10.1038/s41586-018-0030-5

13. Bai F-Y, Han D-Y, Duan S-F, Wang Q-M. The Ecology and Evolution of the Baker’s Yeast Saccharomyces cerevisiae. Genes. 2022;13: 230. doi:10.3390/genes13020230

14. Gallone B, Steensels J, Prahl T, Soriaga L, Saels V, Herrera-Malaver B, et al. Domestication and Divergence of Saccharomyces cerevisiae Beer Yeasts. Cell. 2016;166: 1397–1410.e16. 10.1016/j.cell.2016.08.020

15. Yue J-X, Li J, Aigrain L, Hallin J, Persson K, Oliver K, et al. Contrasting evolutionary genome dynamics between domesticated and wild yeasts. Nature Genetics. 2017;49: 913–924. doi:10.1038/ng.3847

16. Eberlein C, Hénault M, Fijarczyk A, Charron G, Bouvier M, Kohn LM, et al. Hybridization is a recurrent evolutionary stimulus in wild yeast speciation. Nature Communications. 2019;10: 923. doi:10.1038/s41467-019-08809-7

17. Steensels J, Gallone B, Voordeckers K, Verstrepen KJ. Domestication of Industrial Microbes. Current Biology. 2019;29: R381–R393. 10.1016/j.cub.2019.04.025

18. De Chiara M, Barré BP, Persson K, Irizar A, Vischioni C, Khaiwal S, et al. Domestication reprogrammed the budding yeast life cycle. Nature Ecology & Evolution. 2022. doi:10.1038/s41559-022-01671-9

19. Sampaio JP, Pontes A. Yeast domestication. Current Biology. 2025;35: R575–R586. doi:10.1016/j.cub.2025.04.056

20. Bigey F, Segond D, Friedrich A, Guezenec S, Bourgais A, Huyghe L, et al. Evidence for Two Main Domestication Trajectories in Saccharomyces cerevisiae Linked to Distinct Bread-Making Processes. Current Biology. 2021;31: 722–732.e5. doi:10.1016/j.cub.2020.11.016

21. Varela JA, Puricelli M, Ortiz-Merino RA, Giacomobono R, Braun-Galleani S, Wolfe KH, et al. Origin of Lactose Fermentation in Kluyveromyces lactis by Interspecies Transfer of a Neo-functionalized Gene Cluster during Domestication. Current Biology. 2019;29: 4284–4290.e2. doi:10.1016/j.cub.2019.10.044

22. Friedrich A, Gounot J-S, Tsouris A, Bleykasten C, Freel K, Caradec C, et al. Contrasting genomic evolution between domesticated and wild Kluyveromyces lactis yeast populations. Genome Biology and Evolution. 2023 [cited 9 Nov 2023]. doi:10.1093/gbe/evad004

23. Lane MM, Morrissey JP. Kluyveromyces marxianus: A yeast emerging from its sister’s shadow. Fungal Biology Reviews. 2010;24: 17–26. 10.1016/j.fbr.2010.01.001

24. Morrissey JP, Etschmann MMW, Schrader J, de Billerbeck GM. Cell factory applications of the yeast Kluyveromyces marxianus for the biotechnological production of natural flavour and fragrance molecules. Yeast. 2015;32: 3–16. 10.1002/yea.3054

25. Karim A, Gerliani N, Aïder M. Kluyveromyces marxianus: An emerging yeast cell factory for applications in food and biotechnology. International Journal of Food Microbiology. 2020;333: 108818. doi:10.1016/j.ijfoodmicro.2020.108818

26. Rajkumar AS, Morrissey JP. Rational engineering of Kluyveromyces marxianus to create a chassis for the production of aromatic products. Microb Cell Fact. 2020;19: 207. doi:10.1186/s12934-020-01461-7

27. Domenzain I, Sánchez B, Anton M, Kerkhoven EJ, Millán-Oropeza A, Henry C, et al. Reconstruction of a catalogue of genome-scale metabolic models with enzymatic constraints using GECKO 2.0. Nature Communications. 2022;13: 3766. doi:10.1038/s41467-022-31421-1

28. Smets J, Godoy HE, Goossenaerts J, Van Bun E, Deparis Q, Bauwens J, et al. Metabolic engineering and adaptive laboratory evolution of Kluyveromyces marxianus for lactic acid production. Microbial Cell Factories. 2025;24: 179. doi:10.1186/s12934-025-02805-x

29. Zhang M, Ruan Y, Hu Y, Huo Y, Wang M, Zhu Z, et al. A novel high-temperature Kluyveromyces marxianus as a microbial cell factory host for sesquiterpene production. Biochemical Engineering Journal. 2025;220: 109739. doi:10.1016/j.bej.2025.109739

30. Ortiz-Merino RA, Varela JA, Coughlan AY, Hoshida H, da Silveira WB, Wilde C, et al. Ploidy Variation in Kluyveromyces marxianus Separates Dairy and Non-dairy Isolates. Frontiers in Genetics. 2018;9. doi:10.3389/fgene.2018.00094

31. Lozano-Aguirre L, Avitia M, Lappe-Oliveras P, Licona-Cassani C, Cevallos MA, Borgne SL. Draft genomes of four Kluyveromyces marxianus isolates retrieved from the elaboration process of henequen (Agave fourcroydes) mezcal. Microbiology Resource Announcements. 2024. pp. e00861–23. Available: https://journals.asm.org/doi/abs/10.1128/mra.00861-23

32. Moreno-Hernández JM, García-Ortega LF, Morales L, Pohl HC, DeLuna A, Mancera E. Seven draft genomes of Kluyveromyces marxianus strains from alcoholic fermentations in South Africa. Microbiology Resource Announcements. 2026;0: e01042–25. doi:10.1128/mra.01042-25

33. Gómez-Márquez C, Sandoval-Nuñez D, Gschaedler A, Romero-Gutiérrez T, Amaya-Delgado L, Morales JA. Diploid genome assembly of Kluyveromyces marxianus NRRL Y-50883 (SLP1). G3 Genes|Genomes|Genetics. 2021. doi:10.1093/g3journal/jkab347

34. Ibarra-Rivera Gabriel, Muñoz-Miranda Luis Alfonso, Pereira-Santana Alejandro, Arellano-Plaza Melchor, Gschaedler-Mathis Anne, Kirchmayr Manuel R., et al. Draft genome sequence of Kluyveromyces marxianus MCHA. Microbiology Resource Announcements. 2026;0: e01266–25. doi:10.1128/mra.01266-25

35. Gallegos-Casillas P, García-Ortega LF, Espinosa-Cantú A, Avelar-Rivas JA, Torres-Lagunes CG, Cano-Ricardez A, et al. Yeast diversity in open agave fermentations across Mexico. Yeast. 2024;41: 35–51. 10.1002/yea.3913

36. López-Gallegos C, Aguirre-Dugua X, Avelar-Rivas JA, Kirchmayr MR, Morales L, DeLuna A, et al. Ecological divergence of sympatric Saccharomyces species across wild and fermentative environments in the neotropics. bioRxiv. 2025; 2025.05.31.656962. doi:10.1101/2025.05.31.656962

37. Avelar-Rivas JA, Sedeño I, García-Ortega LF, Urban Aragon JA, López-Gallegos C, Aguirre-Dugua X, et al. Recurrent introgression and geographical stratification shape Saccharomyces cerevisiae in the Neotropics. Nature Communications. 2026. doi:10.1038/s41467-026-69138-0

38. Lachance M-A. Phylogenies in yeast species descriptions: In defense of neighbor-joining. Yeast. 2022;39: 513–520. 10.1002/yea.3812

39. Gostincar C. Towards genomic criteria for delineating fungal species. Journal of Fungi. 2020;6: 246.

40. Zonneveld BJM, Steensma HY. Mating, Sporulation and Tetrad Analysis in Kluyveromyces lactis. In: Wolf K, Breunig K, Barth G, editors. Non-Conventional Yeasts in Genetics, Biochemistry and Biotechnology: Practical Protocols. Berlin, Heidelberg: Springer Berlin Heidelberg; 2003. pp. 151–154. doi:10.1007/978-3-642-55758-3_23

41. Boeva V, Popova T, Bleakley K, Chiche P, Cappo J, Schleiermacher G, et al. Control-FREEC: a tool for assessing copy number and allelic content using next-generation sequencing data. Bioinformatics. 2011. pp. 423–425. Available: 10.1093/bioinformatics/btr670

42. Varela JA, Montini N, Scully D, Van der Ploeg R, Oreb M, Boles E, et al. Polymorphisms in the LAC12 gene explain lactose utilisation variability in Kluyveromyces marxianus strains. FEMS Yeast Res. 2017;17. doi:10.1093/femsyr/fox021

43. Gallone B, Mertens S, Gordon JL, Maere S, Verstrepen KJ, Steensels J. Origins, evolution, domestication and diversity of Saccharomyces beer yeasts. Current Opinion in Biotechnology. 2018;49: 148–155. 10.1016/j.copbio.2017.08.005

44. Larralde-Corona CP, De la Torre-González FJ, Vázquez-Landaverde PA, Hahn D, Narváez-Zapata JA. Rational Selection of Mixed Yeasts Starters for Agave Must Fermentation. Frontiers in Sustainable Food Systems. 2021;5. Available: https://www.frontiersin.org/articles/10.3389/fsufs.2021.684228

45. Seehausen O, Butlin RK, Keller I, Wagner CE, Boughman JW, Hohenlohe PA, et al. Genomics and the origin of species. Nature Reviews Genetics. 2014;15: 176–192. doi:10.1038/nrg3644

46. Colón-González M, Aguirre-Dugua X, Guerrero-Osornio MG, Avelar-Rivas JA, DeLuna A, Mancera E, et al. Thriving in Adversity: Yeasts in the Agave Fermentation Environment. Yeast. 2025;42: 16–30. doi:10.1002/yea.3989

47. Jara-Servin A, Alcaraz LD, Juarez-Serrano SI, Espinosa-Jaime A, Barajas I, Morales L, et al. Microbial Communities in Agave Fermentations Vary by Local Biogeographic Regions. Environmental Microbiology Reports. 2025;17: e70057. doi:10.1111/1758-2229.70057

48. Gentry HS. Agaves of Continental North America. University of Arizona Press; 1982. doi:10.2307/j.ctv1t4m2h4

49. Colunga-GarcíaMarín P, Zizumbo-Villarreal D. Tequila and other Agave spirits from west-central Mexico: current germplasm diversity, conservation and origin. Biodiversity and Conservation. 2007;16: 1653–1667. doi:10.1007/s10531-006-9031-z

50. Barsoum E., Rajaei N., Åström S. U. RAS/Cyclic AMP and Transcription Factor Msn2 Regulate Mating and Mating-Type Switching in the Yeast Kluyveromyces lactis. Eukaryotic Cell. 2011;10: 1545–1552. doi:10.1128/ec.05158-11

51. Solieri L, Cassanelli S, Huff F, Barroso L, Branduardi P, Louis EJ, et al. Insights on life cycle and cell identity regulatory circuits for unlocking genetic improvement in Zygosaccharomyces and Kluyveromyces yeasts. FEMS Yeast Research. 2021;21. doi:10.1093/femsyr/foab058

52. Giannakou K, Cotterrell M, Delneri D. Genomic adaptation of Saccharomyces species to industrial environments. Frontiers in Genetics. 2020;Volume 11-2020. doi:10.3389/fgene.2020.00916

53. Christensen KE, Deal A, Wang J-TJ, Duarte A, Edwards JL, Goodman JLN, et al. Signatures of Innovation and Selection in the Extremotolerant Yeast Kluyveromyces marxianus. Genome Biology and Evolution. 2026. Available: 10.1093/gbe/evag060

54. Lachance M-A. Yeast communities in a natural tequila fermentation. Antonie van Leeuwenhoek. 1995;68: 151–160. doi:10.1007/BF00873100

55. Xu Y, Lin Z, Tang C, Tang Y, Cai Y, Zhong H, et al. A new massively parallel nanoball sequencing platform for whole exome research. BMC Bioinformatics. 2019;20: 153. doi:10.1186/s12859-019-2751-3

56. Li J, Zhai Z, Zhang H, Su Z, Liu Y, Chen H, et al. Deep learning enables the use of ultra-high-density array in DNBSEQ. Scientific Reports. 2024;14: 27847. doi:10.1038/s41598-024-78748-x

57. Chen S, Zhou Y, Chen Y, Gu J. fastp: an ultra-fast all-in-one FASTQ preprocessor. Bioinformatics. 2018;34: i884–i890. doi:10.1093/bioinformatics/bty560

58. Prjibelski A, Antipov D, Meleshko D, Lapidus A, Korobeynikov A. Using SPAdes De Novo Assembler. Current Protocols in Bioinformatics. 2020;70: e102. 10.1002/cpbi.102

59. Gurevich A, Saveliev V, Vyahhi N, Tesler G. QUAST: quality assessment tool for genome assemblies. Bioinformatics. 2013;29: 1072–1075. doi:10.1093/bioinformatics/btt086

60. Lertwattanasakul N, Kosaka T, Hosoyama A, Suzuki Y, Rodrussamee N, Matsutani M, et al. Genetic basis of the highly efficient yeast Kluyveromyces marxianus: complete genome sequence and transcriptome analyses. Biotechnology for Biofuels. 2015;8: 47. doi:10.1186/s13068-015-0227-x

61. Kolesov G, Mewes H-W, Frishman D. SNAPping up functionally related genes based on context information: a colinearity-free approach1 1Edited by J. Thornton. Journal of Molecular Biology. 2001;311: 639–656. 10.1006/jmbi.2001.4701

62. Simão FA, Waterhouse RM, Ioannidis P, Kriventseva EV, Zdobnov EM. BUSCO: assessing genome assembly and annotation completeness with single-copy orthologs. Bioinformatics. 2015;31: 3210–3212. doi:10.1093/bioinformatics/btv351

63. The UniProt Consortium. UniProt: a hub for protein information. Nucleic Acids Research. 2015;43: D204–D212. doi:10.1093/nar/gku989

64. Wick R, Menzel P. Filtlong: quality filtering tool for long reads. https://github.com/rrwick/Filtlong. 2019.

65. Koren S, Walenz BP, Berlin K, Miller JR, Bergman NH, Phillippy AM. Canu: scalable and accurate long-read assembly via adaptive k-mer weighting and repeat separation. Genome Research. 2017;27: 722–736. doi:10.1101/gr.215087.116

66. Vaser R, Sović I, Nagarajan N, Šikić M. Fast and accurate de novo genome assembly from long uncorrected reads. Genome Research. 2017;27: 737–746. doi:10.1101/gr.214270.116

67. Li H, Durbin R. Fast and accurate short read alignment with Burrows–Wheeler transform. Bioinformatics. 2009;25: 1754–1760. doi:10.1093/bioinformatics/btp324

68. Inokuma K, Ishii J, Hara KY, Mochizuki M, Hasunuma T, Kondo A. Complete Genome Sequence of Kluyveromyces marxianus NBRC1777, a Nonconventional Thermotolerant Yeast. Genome Announc. 2015;3: e00389–15. doi:doi:10.1128/genomeA.00389-15

69. McKenna A, Hanna M, Banks E, Sivachenko A, Cibulskis K, Kernytsky A, et al. The Genome Analysis Toolkit: A MapReduce framework for analyzing next-generation DNA sequencing data. Genome Research. 2010;20: 1297–1303. doi:10.1101/gr.107524.110

70. Minh BQ, Schmidt HA, Chernomor O, Schrempf D, Woodhams MD, von Haeseler A, et al. IQ-TREE 2: New Models and Efficient Methods for Phylogenetic Inference in the Genomic Era. Molecular Biology and Evolution. 2020;37: 1530–1534. doi:10.1093/molbev/msaa015

71. Danecek P, Auton A, Abecasis G, Albers CA, Banks E, DePristo MA, et al. The variant call format and VCFtools. Bioinformatics. 2011;27: 2156–2158. doi:10.1093/bioinformatics/btr330

72. Zheng X, Levine D, Shen J, Gogarten SM, Laurie C, Weir BS. A high-performance computing toolset for relatedness and principal component analysis of SNP data. Bioinformatics. 2012;28: 3326–3328. doi:10.1093/bioinformatics/bts606

73. Alexander DH, Novembre J, Lange K. Fast model-based estimation of ancestry in unrelated individuals. Genome Research. 2009;19: 1655–1664. doi:10.1101/gr.094052.109

74. Purcell S, Neale B, Todd-Brown K, Thomas L, Ferreira MAR, Bender D, et al. PLINK: A tool set for whole-genome association and population-based linkage analyses. American Journal of Human Genetics. 2007;81: 559–575. doi:10.1086/519795

75. Behr AA, Liu KZ, Liu-Fang G, Nakka P, Ramachandran S. pong: fast analysis and visualization of latent clusters in population genetic data. Bioinformatics. 2016;32: 2817–2823. doi:10.1093/bioinformatics/btw327

76. Francis RM. pophelper: an R package and web app to analyse and visualize population structure. Molecular Ecology Resources. 2017;17: 27–32. 10.1111/1755-0998.12509

77. Jain C, Rodriguez-R LM, Phillippy AM, Konstantinidis KT, Aluru S. High throughput ANI analysis of 90K prokaryotic genomes reveals clear species boundaries. Nature Communications. 2018;9: 5114. doi:10.1038/s41467-018-07641-9

78. Emms DM, Kelly S. OrthoFinder: phylogenetic orthology inference for comparative genomics. Genome Biology. 2019;20: 238. doi:10.1186/s13059-019-1832-y

79. Marcais G, Delcher AL, Phillippy AM, Coston R, Salzberg SL, Zimin A. MUMmer4: A fast and versatile genome alignment system. PLOS Computational Biology. 2018;14: e1005944. doi:10.1371/journal.pcbi.1005944

80. Tang H, Krishnakumar V, Zeng X, Xu Z, Taranto A, Lomas JS, et al. JCVI: A versatile toolkit for comparative genomics analysis. iMeta. 2024;3: e211. doi:10.1002/imt2.211

81. Tellini N, De Chiara M, Mozzachiodi S, Tattini L, Vischioni C, Naumova ES, et al. Ancient and recent origins of shared polymorphisms in yeast. Nature Ecology & Evolution. 2024;8: 761–776. doi:10.1038/s41559-024-02352-5

82. Petr M, Vernot B, Kelso J. admixr—R package for reproducible analyses using ADMIXTOOLS. Bioinformatics. 2019;35: 3194–3195. doi:10.1093/bioinformatics/btz030

83. Steenwyk JL, Opulente DA, Kominek J, Shen X-X, Zhou X, Labella AL, et al. Extensive loss of cell-cycle and DNA repair genes in an ancient lineage of bipolar budding yeasts. PLOS Biology. 2019;17: e3000255. doi:10.1371/journal.pbio.3000255

84. Alexa A, Rahnenfuhrer J. topGO: Enrichment Analysis for Gene Ontology, R package version 2.62.0. 2024. 2024. 10.18129/B9.bioc.topGO

85. Sayols S. rrvgo: a Bioconductor package for interpreting lists of Gene Ontology terms. 2023. doi:10.17912/MICROPUB.BIOLOGY.000811

86. Rosebrock AP. Analysis of the Budding Yeast Cell Cycle by Flow Cytometry. Cold Spring Harbor Protocols. 2017;2017: pdb.prot088740. doi:10.1101/pdb.prot088740

87. Kassir Y, Simchen G. Monitoring meiosis and sporulation in Saccharomyces cerevisiae. Methods in Enzymology. Academic Press; 1991. pp. 94–110. doi:10.1016/0076-6879(91)94009-2

88. Guthrie C, Fink GR. Guide to yeast genetics and Molecular Biology. Methods in Enzymology. Academic Press; 1991. p. xvii. doi:10.1016/0076-6879(91)94001-S

89. Wu L, Wang M, Zha G, Zhou J, Yu Y, Lu H. A protocol of rapid laboratory evolution by genome shuffling in Kluyveromyces marxianus. MethodsX. 2020;7: 101138. 10.1016/j.mex.2020.101138

90. Mancera E, Frazer C, Porman AM, Ruiz-Castro S, Johnson AD, Bennett RJ. Genetic modification of closely related Candida species. Frontiers in Microbiology. 2019;10: 357.

91. Kooistra RA, Steensma HY. Transformation of Kluyveromyces lactis. In: Wolf K, Breunig K, Barth G, editors. Non-Conventional Yeasts in Genetics, Biochemistry and Biotechnology: Practical Protocols. Berlin, Heidelberg: Springer Berlin Heidelberg; 2003. pp. 169–174. doi:10.1007/978-3-642-55758-3_26

92. Goldstein AL, McCusker JH. Three new dominant drug resistance cassettes for gene disruption in Saccharomyces cerevisiae. Yeast. 1999;15: 1541–1553. 10.1002/(SICI)1097-0061(199910)15:14<1541::AID-YEA476>3.0.CO;2-K

93. Ongay-Larios L, Navarro-Olmos R, Kawasaki L, Velázquez-Zavala N, Sánchez-Paredes E, Torres-Quiroz F, et al. Kluyveromyces lactis sexual pheromones. Gene structures and cellular responses to α-factor. FEMS Yeast Research. 2007;7: 740–747. doi:10.1111/j.1567-1364.2007.00249.x

94. Mancilla-Margalli NA, López MG. Generation of Maillard Compounds from Inulin during the Thermal Processing of Agave tequilana Weber Var. azul. Journal of Agricultural and Food Chemistry. 2002;50: 806–812. doi:10.1021/jf0110295

95. Espinosa-Cantú A, Ascencio D, Herrera-Basurto S, Xu J, Roguev A, Krogan NJ, et al. Protein Moonlighting Revealed by Noncatalytic Phenotypes of Yeast Enzymes. Genetics. 2018. pp. 419–431. Available: 10.1534/genetics.117.300377

96. Gu Z, Eils R, Schlesner M. Complex heatmaps reveal patterns and correlations in multidimensional genomic data. Bioinformatics. 2016;32: 2847–2849. doi:10.1093/bioinformatics/btw313

97. Patil I. Visualizations with statistical details: The’ggstatsplot’approach. Journal of Open Source Software. 2021;6: 3167.

