## Supplementary Figures for "Domestication and deep lineage divergence define two discrete trajectories in *Kluyveromyces marxianus*"

### Supplementary Material

#### Supplementary Tables

**Table S1.** Metadata of *K. marxianus* strains included in this study. (XLS file)

**Table S2.** ITS-D1/D2 sequence identity of the *K. marxianus* strains from the three lineages. (XLS file)

**Table S3.** Sporulation efficiency of diploid strains under different induction conditions. (XLS file)

**Table S4.** Patterson's D statistics to test gene flow between clades and subpopulations. (XLS file)

**Table S5.** Gene Ontology enrichment analysis of clade B-specific gene losses. (XLS file)

**Table S6.** Genes with CNVs detected across the *K. marxianus* collection. (XLS file)

**Table S7.** Conditions used for high-throughput phenotyping. (XLS file)

#### Supplementary Figures

**Figure S1.** Population diversity estimates across 178 *K. marxianus* strains

**Figure S2.** Population structure of *K. marxianus* inferred from genome-wide SNPs

**Figure S3.** Geographic origin and phylogenetic position of *K. marxianus* strains from Mexico

**Figure S4.** Ploidy variation and aneuploidy in *K. marxianus* genomes

**Figure S5.** BUSCO assessment of *K. marxianus* genome annotations

**Figure S6.** Assessment of reciprocal monophyly of Clade C

**Figure S7.** Conserved synteny across genomes from the three lineages

**Figure S8.** Low rates of gene flow suggest reproductive isolation of Clade C

**Figure S9.** Gene copy-number variation patterns across the *K. marxianus* genomes

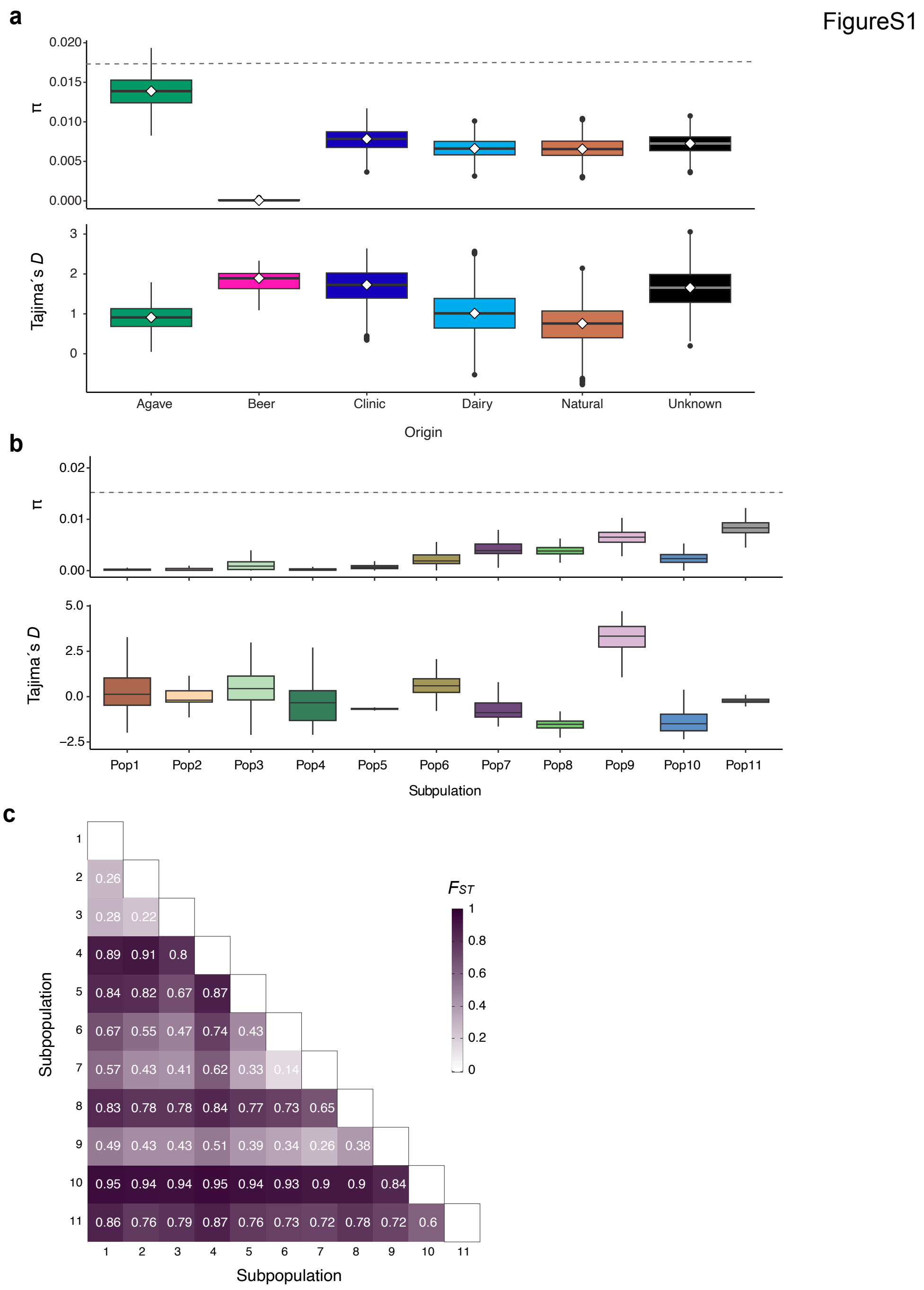


**Figure S1. Population diversity and differentiation estimates across 178 *K. marxianus strains*. a)** Nucleotide diversity (π) and Tajima's *D* for strains grouped by isolation origin and **b)** for the eleven subpopulations inferred with Admixture. The dotted gray line indicates the overall species-wide nucleotide diversity ($\pi$=1.6x$10^{-2}$). **c)** Pairwise genetic differentiation ($F_{ST}$) among the eleven subpopulations, estimated using the Weir and Cockerham method. Higher values indicate greater differentiation between subpopulations.

FigureS2

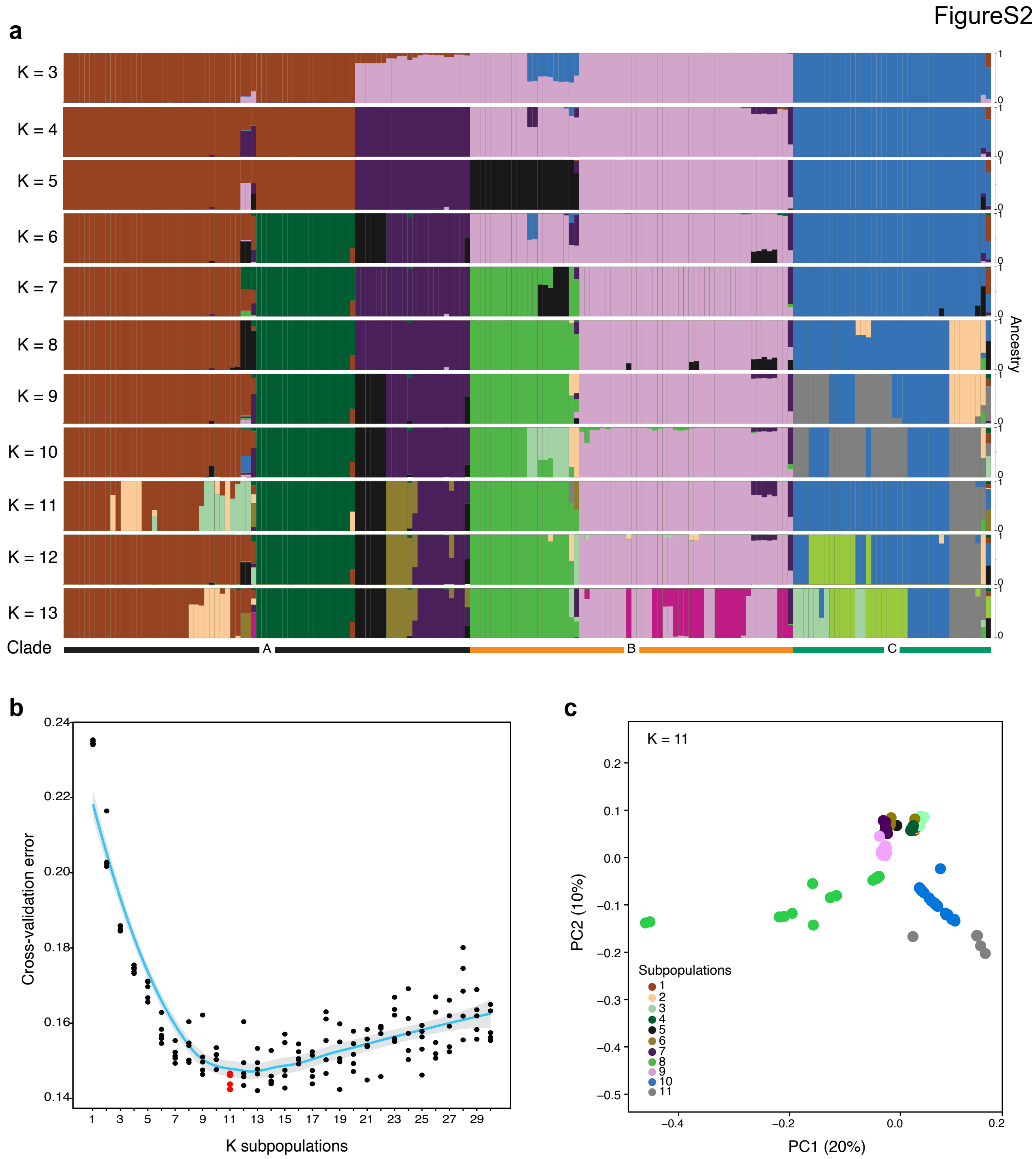


**Figure S2. Population structure of *K. marxianus* inferred from genome-wide SNPs. a)** Admixture results showing ancestry proportions for 178 strains across K=3 to K=13 inferred ancestral subpopulations. Each vertical bar represents an individual genome, with colors indicating inferred ancestry components. Labels at the bottom indicate the higher-level clade assignments A, B, and C, defined from phylogenomic analyses. **b)** Cross-validation error across K values from 1 to 30, used to identify the best-supported number of ancestral subpopulations. The cyan line represents the fitted trend in cross-validation error. The lowest cross-validation error was observed at K=11, highlighted in red. **c)** Principal component analysis of the genome-wide SNP dataset, showing the projection of samples along PC1 and PC2. Each point represents a genome, colored according to its subpopulation assignment at K=11. The PCA recapitulates the structure inferred by Admixture, showing clear differentiation among the major genetic groups.

FigureS3

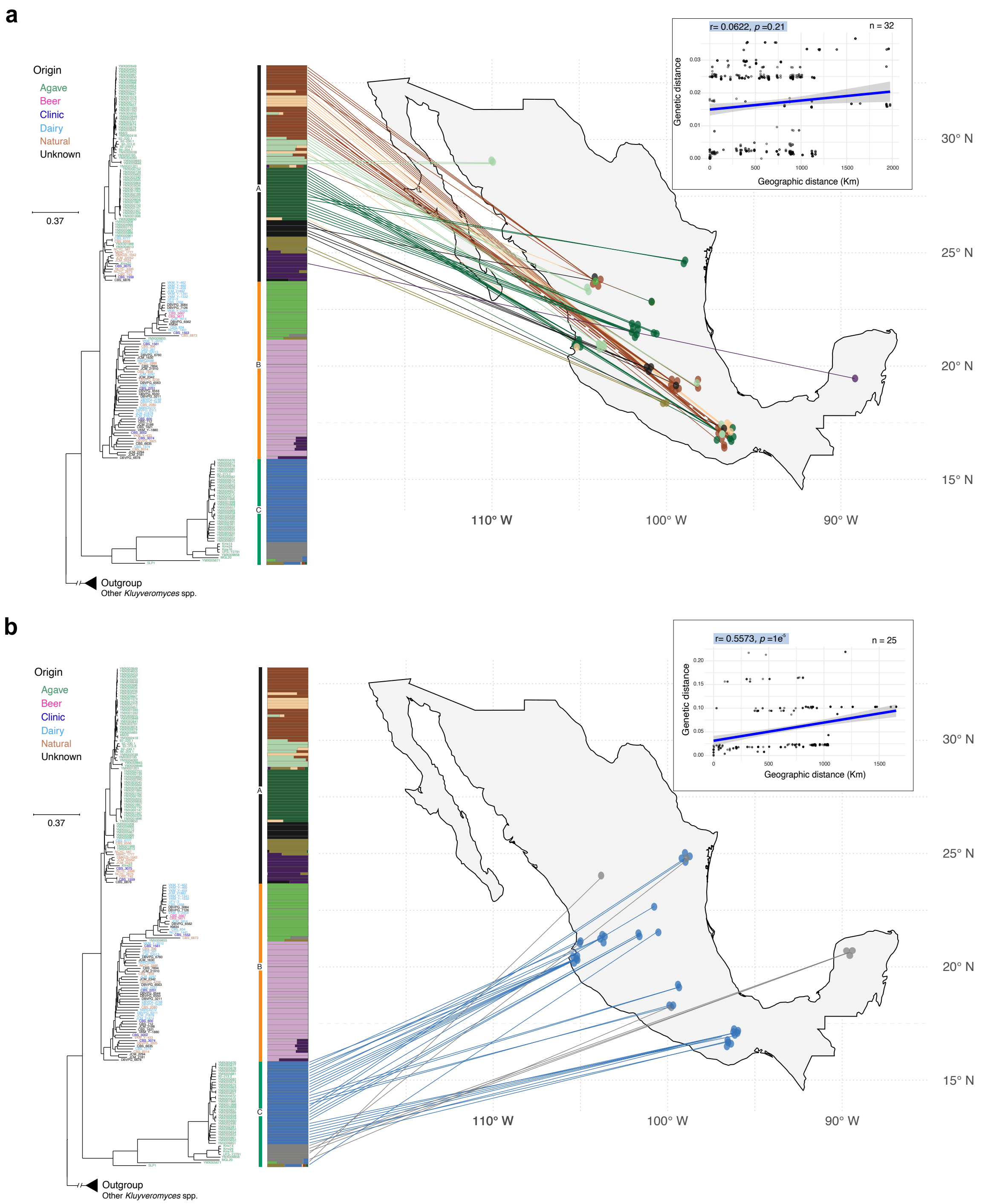


**Figure S3. Geographic origin and phylogenetic position of *K. marxianus* strains from Mexico.** Projection of the geographic distribution of Clade A (**a**) and Clade C (**b**) strains according to their phylogenetic placement. Inset plots for each panel show the correlation between geographic and genetic distances for 32 Clade A strains and 25 Clade C isolates according to the Mantel test. The correlation coefficient and the associated significance (*p*-value), estimated after 10,000 iterations, are highlighted in blue.

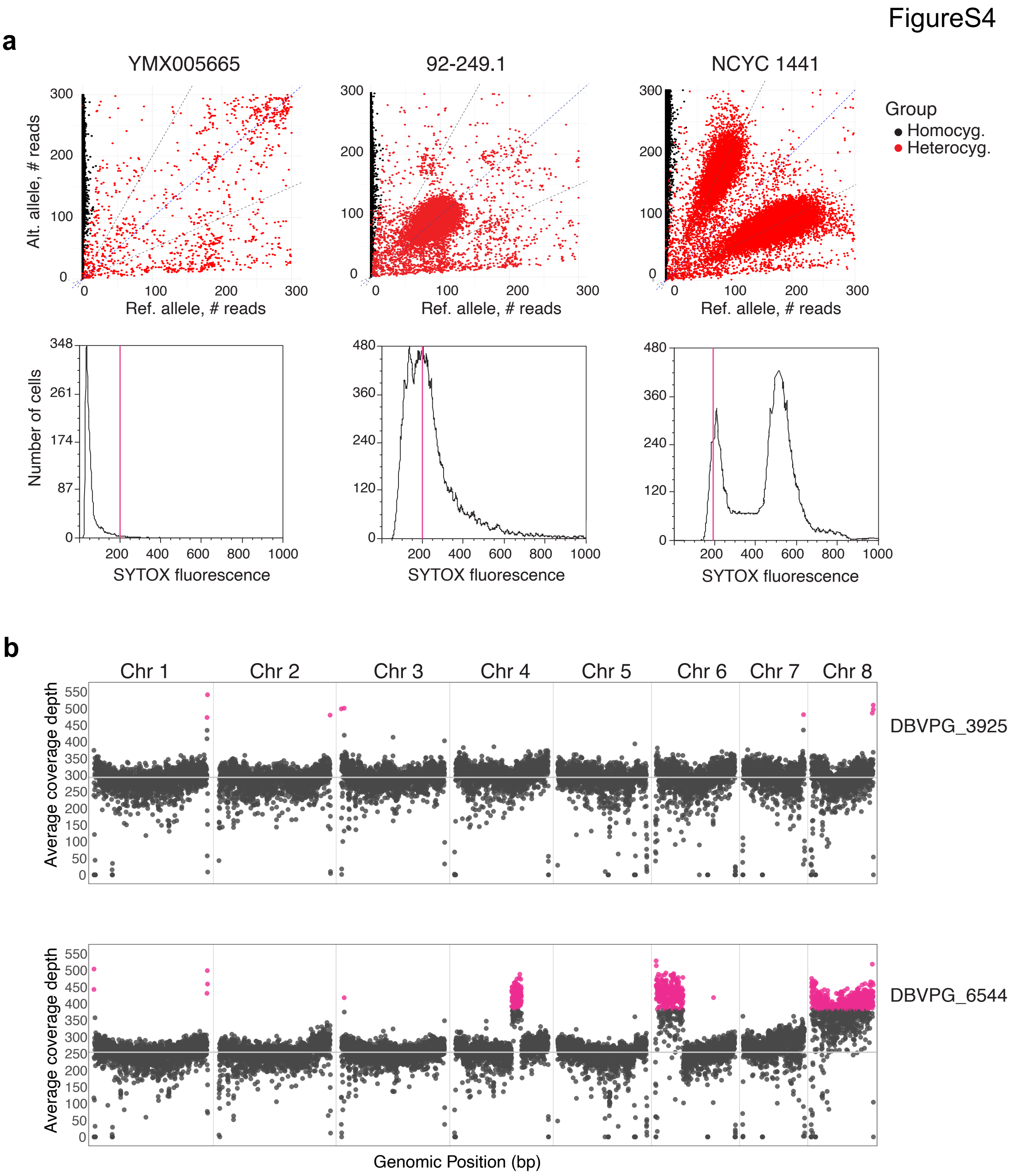


**Figure S4. Ploidy variation and aneuploidy in *K. marxianus* genomes. a)** Examples of *in silico* ploidy inference based on allele-frequency distributions and flow cytometry. Top panels show allele-frequency scatterplots comparing read counts for alternative and reference alleles in strains YMX005665, 92-249.1, and NCYC 1441, representing haploid, diploid, and polyploid genomes, respectively. Bottom panels show the flow cytometry histograms of SYTOX fluorescence intensity for the same strains, reflecting cellular DNA content and supporting the inferred ploidy differences. **b)** Genome-wide coverage depth across the eight *K. marxianus* chromosomes in representative euploid and aneuploid genomes. DBVPG 3925 shows a euploid coverage profile, whereas DBVPG 6544 shows chromosome-level coverage deviations consistent with aneuploidy. Both strains belong to clade B, but DBVPG 6544 (bottom) was specifically isolated from an industrial environment. Aneuploidies were detected from chromosome-level normalized coverage calculated in 1 kb windows. Windows with coverage deviations consistent with aneuploidy are highlighted as magenta points.

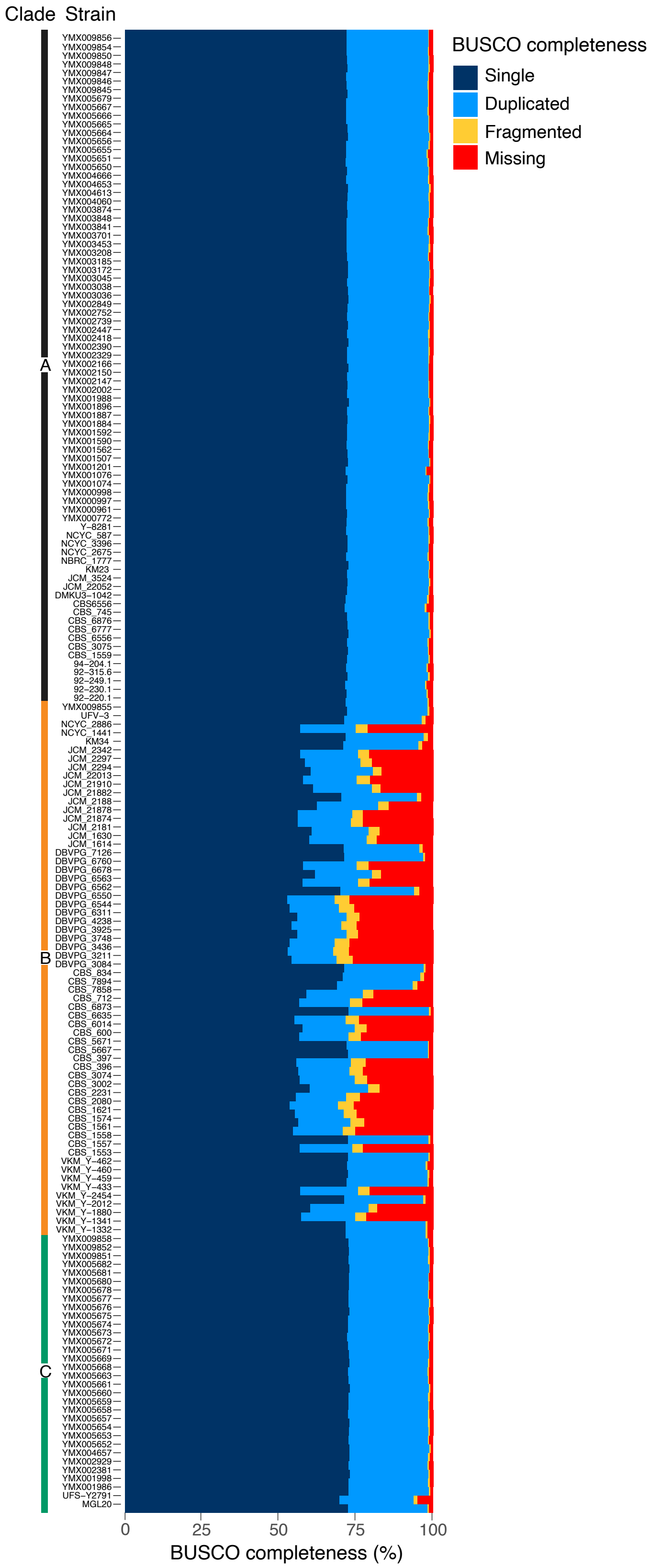
Clade
Strain
A
B
C
YMX009856
YMX009854
YMX009850
YMX009848
YMX009847
YMX009846
YMX009845
YMX005679
YMX005667
YMX005666
YMX005665
YMX005664
YMX005656
YMX005655
YMX005651
YMX005650
YMX004666
YMX004653
YMX004613
YMX004060
YMX003874
YMX003848
YMX003841
YMX003701
YMX003453
YMX003208
YMX003185
YMX003172
YMX003045
YMX003038
YMX003036
YMX002849
YMX002752
YMX002739
YMX002447
YMX002418
YMX002390
YMX002329
YMX002166
YMX002150
YMX002147
YMX002002
YMX001988
YMX001896
YMX001887
YMX001884
YMX001592
YMX001590
YMX001562
YMX001507
YMX001201
YMX001076
YMX001074
YMX000998
YMX000997
YMX000961
YMX000772
Y−8281
NCYC_587
NCYC_3396
NCYC_2675
NBRC_1777
KM23
JCM_3524
JCM_22052
DMKU3−1042
CBS6556
CBS_745
CBS_6876
CBS_6777
CBS_6556
CBS_3075
CBS_1559
94−204.1
92−315.6
92−249.1
92−230.1
92−220.1
YMX009855
UFV−3
NCYC_2886
NCYC_1441
KM34
JCM_2342
JCM_2297
JCM_2294
JCM_22013
JCM_21910
JCM_21882
JCM_2188
JCM_21878
JCM_21874
JCM_2181
JCM_1630
JCM_1614
DBVPG_7126
DBVPG_6760
DBVPG_6678
DBVPG_6563
DBVPG_6562
DBVPG_6550
DBVPG_6544
DBVPG_6311
DBVPG_4238
DBVPG_3925
DBVPG_3748
DBVPG_3436
DBVPG_3211
DBVPG_3084
CBS_834
CBS_7894
CBS_7858
CBS_712
CBS_6873
CBS_6635
CBS_6014
CBS_600
CBS_5671
CBS_5667
CBS_397
CBS_396
CBS_3074
CBS_3002
CBS_2231
CBS_2080
CBS_1621
CBS_1574
CBS_1561
CBS_1558
CBS_1557
CBS_1553
VKM_Y−462
VKM_Y−460
VKM_Y−459
VKM_Y−433
VKM_Y−2454
VKM_Y−2012
VKM_Y−1880
VKM_Y−1341
VKM_Y−1332
YMX009858
YMX009852
YMX009851
YMX005682
YMX005681
YMX005680
YMX005678
YMX005677
YMX005676
YMX005675
YMX005674
YMX005673
YMX005672
YMX005671
YMX005669
YMX005668
YMX005663
YMX005661
YMX005660
YMX005659
YMX005658
YMX005657
YMX005654
YMX005653
YMX005652
YMX004657
YMX002929
YMX002381
YMX001998
YMX001986
UFS−Y2791
MGL20
0
25
50
75
100
BUSCO completeness (%)
BUSCO completeness
Single
Duplicated
Fragmented
Missing

FigureS5

**Figure S5. BUSCO assessment of *K. marxianus* genome annotations.** Stacked bar plot summarizing BUSCO completeness for each annotated genome. Genome assemblies are sorted according to the phylogeny in Figure 1b and grouped by clade. Bars indicate the proportion of BUSCO genes classified as complete, fragmented, or missing as shown in the legend. Most assemblies from clades A and C show high BUSCO completeness, whereas several clade B assemblies show higher proportions of fragmented and missing BUSCOs, consistent with the broader pattern of gene-content reduction observed in this lineage.

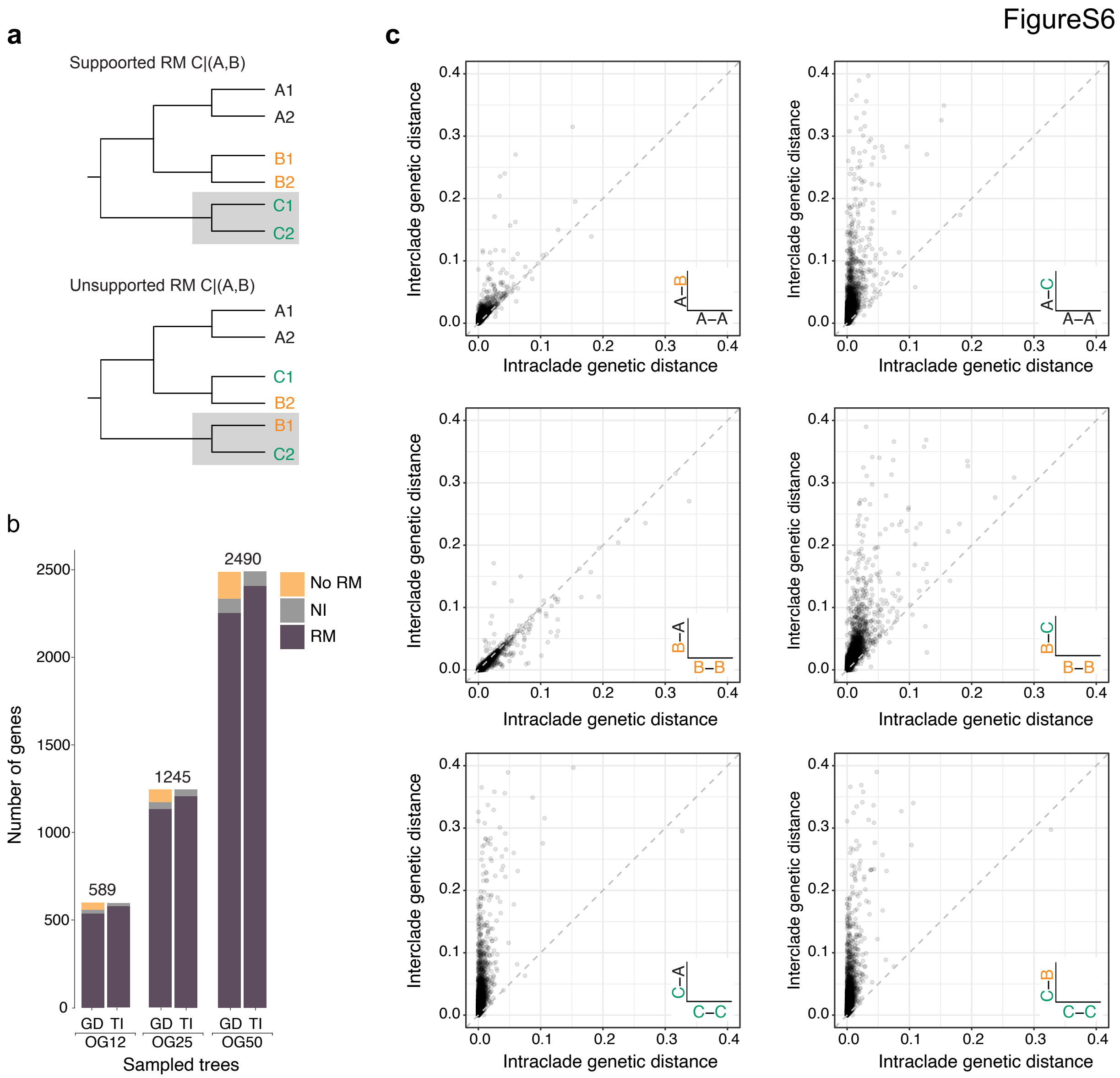


**Figure S6. Assessment of reciprocal monophyly of Clade C. a)** Schematic cladograms illustrating gene-tree topologies that support reciprocal monophyly of clade C (upper tree) or fail to support it (lower tree). **b)** Evaluation of reciprocal monophyly across gene-tree subsets using two complementary approaches: a genetic-distance criterion (GD) and topology inspection (TI). Analyses were performed on three randomly sampled orthogroup datasets of increasing size: OG12 (589 genes), OG25 (1,245 genes), and OG50 (2,490 genes). **c)** Scatterplots comparing within-clade and between-clade genetic distances across gene trees. Points near the diagonal indicate similar within- and between-clade distances. This pattern is maintained for comparisons involving clades A and B, whereas comparisons involving clade C show stronger separation between within- and between-clade distances, consistent with reduced within-clade divergence relative to divergence from other clades and supporting reciprocal monophyly of clade C.

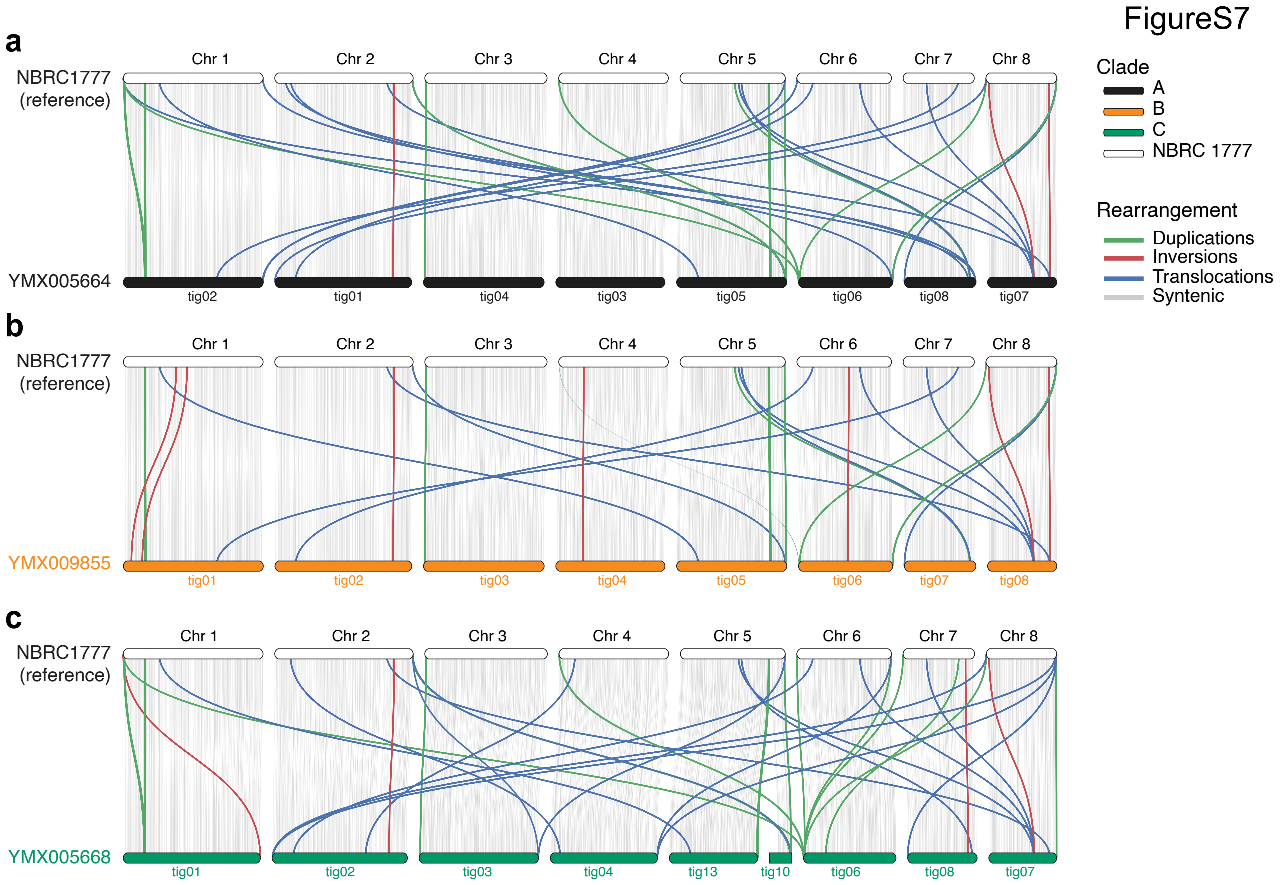


**Figure S7. Conserved synteny across genomes from the three lineages.** Synteny analysis of near-chromosome-level de novo assemblies representing the three main *K. marxianus* lineages. Genomes from strains **a)** YMX005664, **b)** YMX009855, and **c)** YMX005668 were generated using Oxford Nanopore long-read sequencing. Reference sequences from NBRC1777 were aligned against each assembly using BLAST to identify conserved syntenic regions and structural rearrangements. Gray links indicate syntenic regions; red, blue, and green links indicate inversions, duplications, and translocations, respectively.

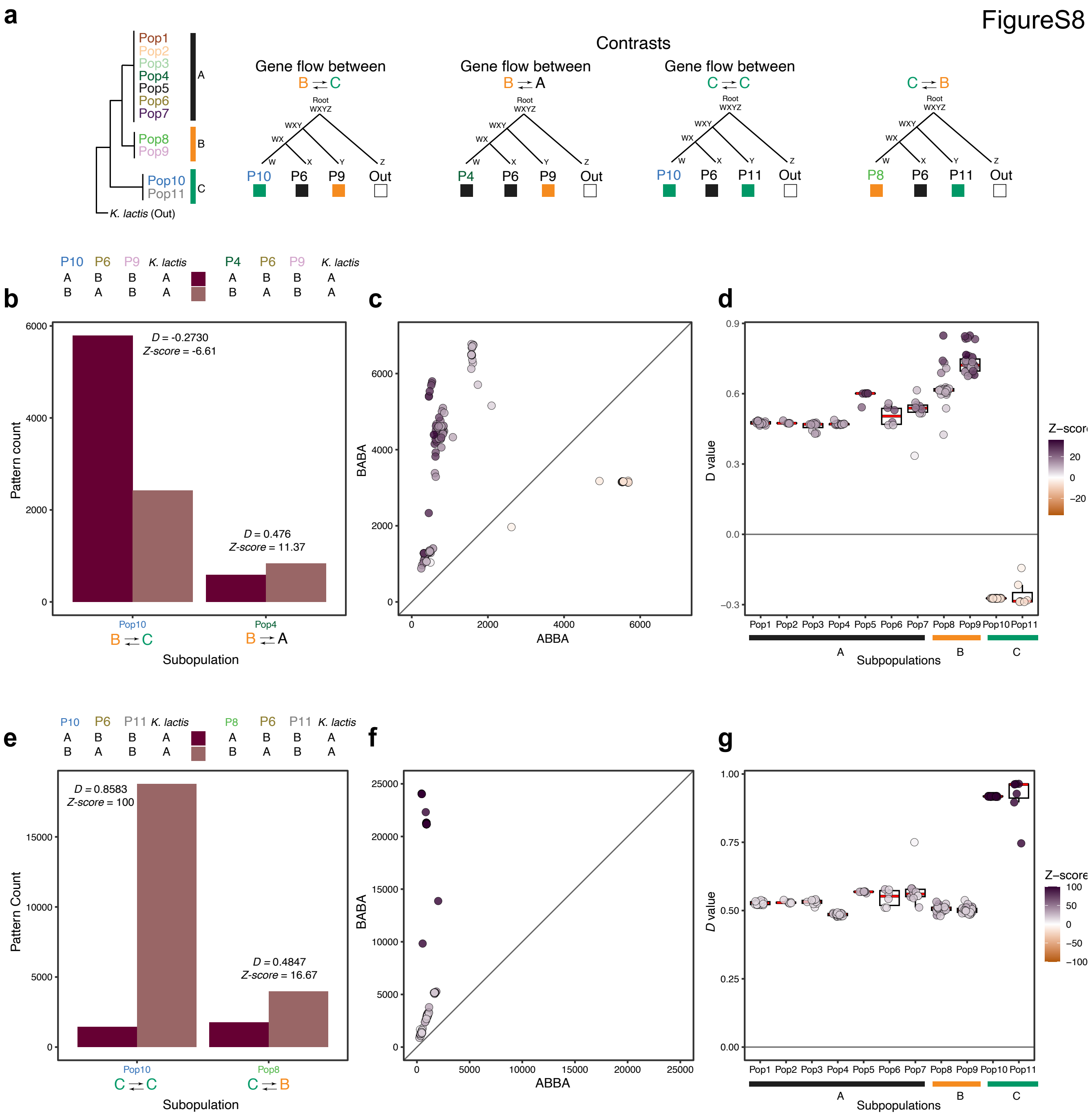


**Figure S8. Low rates of gene flow suggest reproductive isolation of clade C. a)** Patterson's *D*-statistic contrasts used to test for gene flow among the eleven Admixture subpopulations. The cladogram shows their phylogenetic relationships within the three major clades. Each contrast defines a target subpopulation (W), a comparison subpopulation (*X*), a reference subpopulation (*Y*), and *K. lactis* CBS 683 as the outgroup (*Z*). Subpopulation colors match the Admixture assignments in Figure 1; boxes below each subpopulation indicate clade membership. **b)** ABBA-BABA block counts for inter-clade contrasts involving Pop10 from clade C and Pop4 from clade A as target subpopulations, using Pop9 from clade B as the reference. **c)** Scatter plot of ABBA and BABA counts across *K.*

*marxianus* subpopulations using Pop9 as reference. The dotted diagonal indicates equal support for ABBA and BABA patterns, expected in the absence of excess allele sharing. **d)** Distribution of Patterson's *D* values across the eleven subpopulations for the contrasts shown in panel c. **e–g)** Equivalent analyses comparing intra-clade C and inter-clade contrasts using Pop11 from clade C as the reference subpopulation.

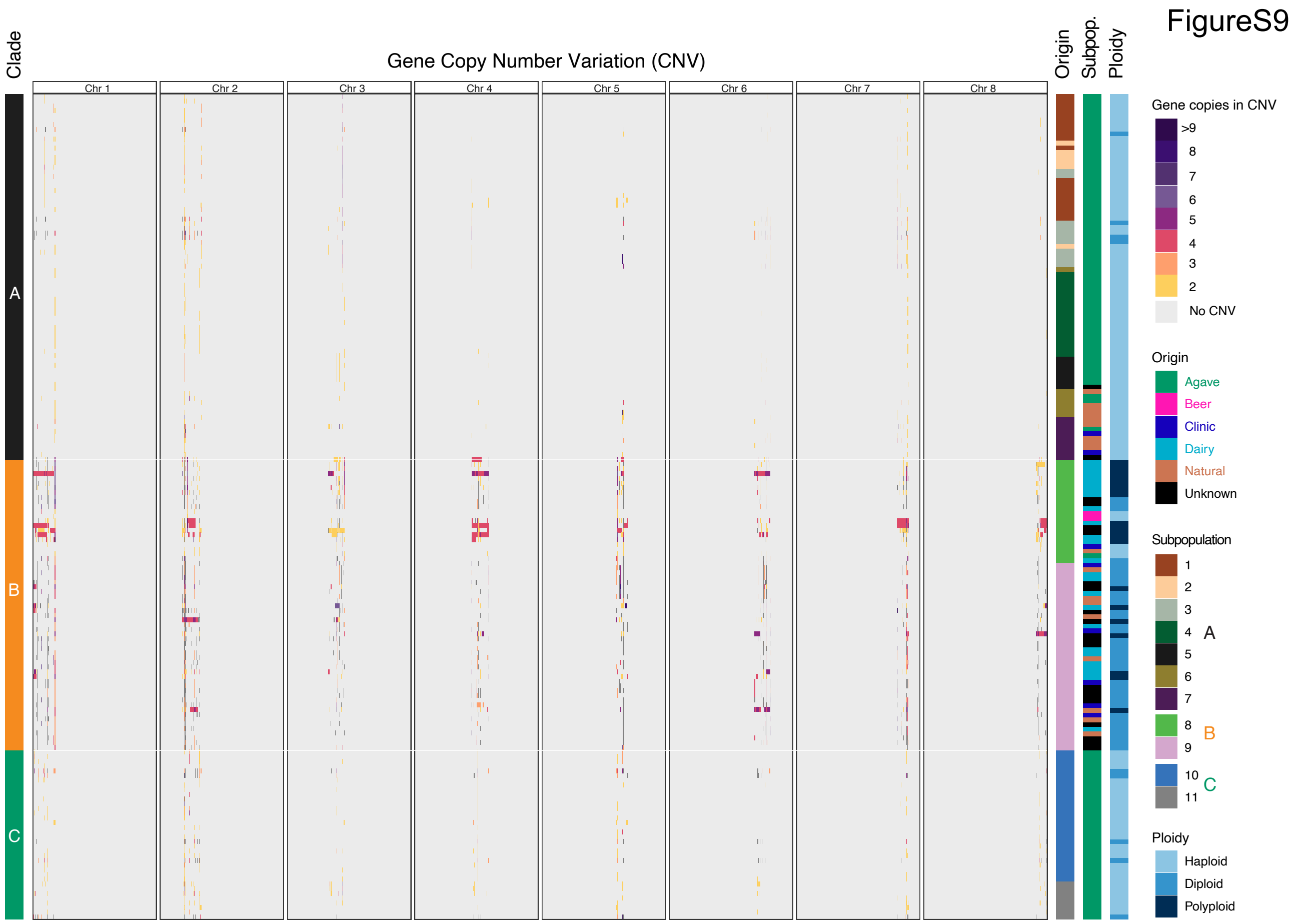


**Figure S9. Gene copy-number variation patterns across the *K. marxianus* genomes**. Heatmap showing gene-level CNV events across 178 genomes from the three phylogenetic clades. Rows represent individual genomes ordered by Admixture profile, and columns represent genes with at least one CNV event, ordered by chromosomal position ($n$ = 4,173 genes). Colors indicate inferred copy number, as shown in the legend. The annotation bar on the left indicates the main phylogenetic clade of each strain, and annotation bars on the right indicate isolation origin, Admixture subpopulation, and ploidy.
